# A family of small proteins links c-di-GMP riboswitch signaling with sporulation in *Clostridioides difficile*

**DOI:** 10.64898/2026.06.10.731418

**Authors:** Adriana Badilla Lobo, Anne-Judith Waligora-Dupriet, Louise Penven, Roland Seifert, César Rodríguez, Frédéric Barbut, Olga Soutourina, Johann Peltier

## Abstract

Cyclic diguanosine monophosphate (c-di-GMP) is a ubiquitous bacterial second messenger that coordinates lifestyle transitions, virulence, and developmental processes. In *Clostridioides difficile*, elevated c-di-GMP levels inhibit sporulation, but the underlying mechanism remained unclear. Here, we identified a conserved family of small, membrane-associated proteins encoded by c-di-GMP riboswitch-regulated genes. Transcriptomic analyses revealed that c-di-GMP represses these genes, and reporter assays demonstrated riboswitch-dependent transcriptional regulation via premature termination mechanism.

Overexpression of a single member, *CD1980.2*, was sufficient to trigger the transcriptional activation of sporulation genes, including sigma factors and their regulons, and to increase spore formation. Conversely, sporulation efficiency decreased proportionally with the number of deleted small protein genes, and the strain lacking all seven genes displayed a severe sporulation defect, underscoring their cumulative and functionally redundant roles. Elevating c-di-GMP levels in the deletion mutant did not further reduce sporulation, supporting a model in which c-di-GMP inhibits spore formation by repressing the expression of this small-protein family. Our work establishes these small proteins, encoded by c-di-GMP riboswitch-regulated genes, as important mediators of developmental output in *C. difficile*. Their redundancy, conservation, and integration into riboswitch regulatory pathways highlight their central role in spore formation, a process essential for pathogen persistence and transmission. These findings expand the repertoire of components regulated by second-messenger signaling in bacterial physiology.

## INTRODUCTION

Bacteria rely on intracellular signaling molecules to rapidly adapt their physiology to fluctuating environments. Among these signals, cyclic diguanosine monophosphate (c-di-GMP) is one of the most widely distributed and well-characterized second messengers ^1,2^. It coordinates numerous processes by modulating the activity of receptor proteins, transcription factors, and RNA-based regulators such as riboswitches ^2,3^. The primary and conserved function of c-di-GMP is to control the transition between motile planktonic growth and sessile community-associated lifestyles, including biofilms ^1,4,5^. Beyond this central role, c-di-GMP is also involved in virulence, cell cycle progression, and developmental pathways ^1,6^. Cellular levels of c-di-GMP depend on the activity of two families of antagonistic enzymes; the diguanylate cyclases (DGCs) which synthesize c-di-GMP and the phosphodiesterases (PDEs), which degrade it ^7–9^.

In *Clostridioides difficile*, a major nosocomial pathogen responsible for antibiotic-associated diarrhea and life-threatening colitis ^10^, c-di-GMP plays a central role in regulating multiple aspects of bacterial physiology ^11,12^. The strain 630 encodes 37 proteins related to c-di-GMP turnover, including 15 predicted active diguanylate cyclases (DGCs) and 18 phosphodiesterases (PDEs), suggesting a tight control of intracellular c-di-GMP levels ^13^. *C. difficile* predominantly relies on c-di-GMP-responsive riboswitches to regulate gene expression, with 12 predicted class I and 4 class II riboswitches in strain 630 ^14–17^. Class I riboswitches generally act as “OFF” switches, whereas class II riboswitches function as “ON” switches ^14,18^. High c-di-GMP concentrations inhibit flagellar gene expression and motility while promoting adhesion, aggregation, and biofilm formation ^19–25^. For example, the class I riboswitch Cdi1_3 represses the early flagellar operon, whereas class II riboswitches (Cdi2_1, Cdi2_3, Cdi2_4) activate expression of surface adhesins CD2831, CD3246, and type IV pili ^14,21,24,26^. Conversely, low c-di-GMP levels activate the metalloprotease PPEP-1 through the riboswitch Cdi1_12, which cleaves CD2831and CD3246 from the cell surface, promoting dispersal ^23,24,27^. Additional regulation occurs through the *cmrRST* operon, whose expression is activated by a class II c-di-GMP riboswitch (Cdi2_2). This operon encodes a signal transduction system, which modulates motility, biofilm formation, and colony morphology^28,29^.

Beyond lifestyle transitions, c-di-GMP influences virulence and stress adaptation in *C. difficile*. It represses the expression of the flagellar sigma factor σ^D^, indirectly reducing toxin production ^19,30,31^, and regulates prophage-encoded type I toxin-antitoxin systems through dedicated riboswitches (Cdi1_4 and Cdi1_5) ^32^. Importantly, c-di-GMP also regulates sporulation in *C. difficile*. Elevated c-di-GMP inhibits early sporulation gene expression, whereas reducing c-di-GMP via PDE PdcB overexpression enhances sporulation ^33,34^.

However, the molecular effectors linking c-di-GMP to sporulation, the key developmental process that enables *C. difficile* persistence and transmission ^35,36^, remained unknown.

Sporulation is a tightly regulated developmental process triggered by environmental cues such as nutrient limitation, population density, and possibly other unidentified signals ^37^.

Activation of the master regulator Spo0A induces asymmetric cell division, producing a forespore and a mother cell ^38,39^. Gene expression in these compartments is sequentially controlled by four cell type-specific sigma factors that are sequentially activated ^35,40^. The sigma factors σ^F^ and σ^E^ direct early stages in the forespore and mother cell, respectively, and are subsequently replaced by σ^G^ and σ^K^ following forespore engulfment. While the overall architecture of this regulatory cascade resembles that of *Bacillus subtilis*, several key differences exist ^41,42^.

In this study, we report the identification of a conserved family of small, membrane-associated proteins, encoded by c-di-GMP riboswitch-regulated genes, linking c-di-GMP signaling to sporulation in *C. difficile*. Transcriptomic analyses reveal that these proteins are repressed by c-di-GMP via class I riboswitches. Functional studies demonstrate that their overexpression promotes sporulation, while gene deletion severely reduces spore formation. Together, these findings expand the repertoire of genes regulated by c-di-GMP-responsive riboswitches and uncover a previously unrecognized mechanism by which this second messenger controls sporulation in *C. difficile*.

## RESULTS

### C-di-GMP alters riboswitch-controlled gene expression and represses sporulation-related genes in *C. difficile*

To elevate intracellular c-di-GMP levels we integrated the inducible *P_tet_* promoter upstream of the diguanylate cyclase gene *dccA* in *C. difficile* 630Δ*erm*, a commonly used spontaneous erythromycin-sensitive derivative of the strain 630. Upon anydrotetracycline (ATc) induction, the resulting strain, *P_tet_-dccA*, displayed a marked increase in intracellular c-di-GMP concentration in comparison with wild type as confirmed by LC-MS/MS (Fig. 1A).

**Fig. 1.**
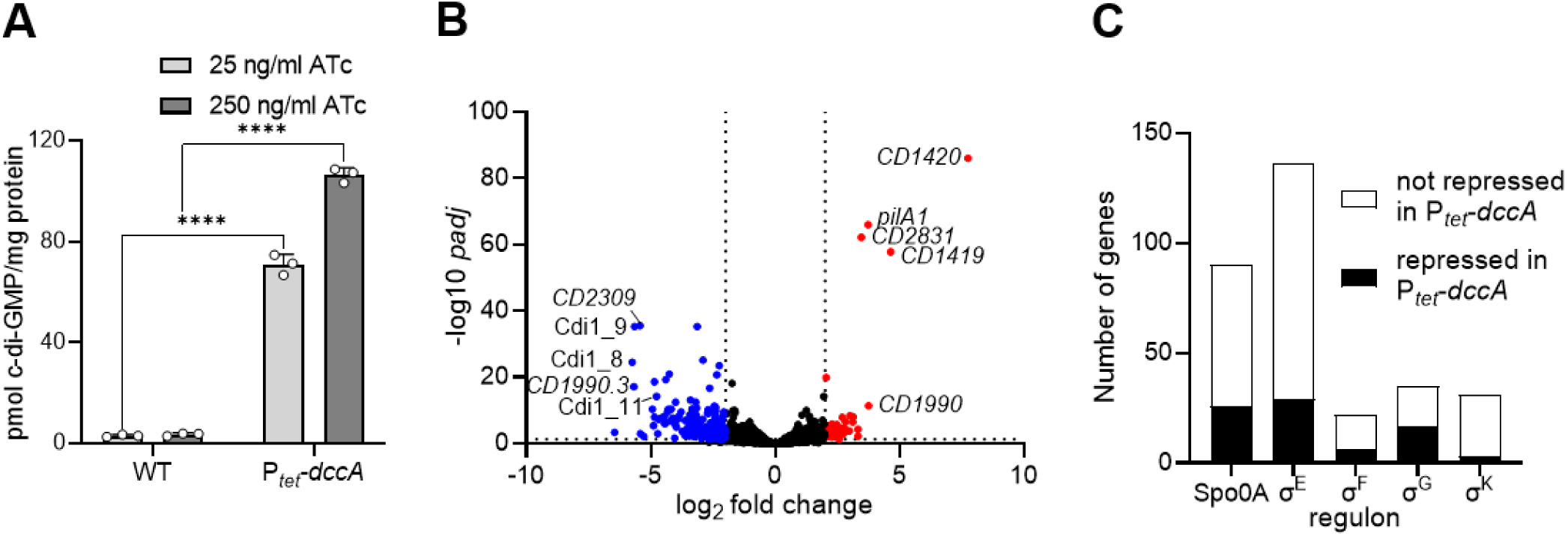
Elevated intracellular c-di-GMP concentration modulates expression of c-di-GMP-responsive loci and represses expression of sporulation genes in *C. difficile*. (A) Intracellular c-di-GMP concentrations in *C. difficile* 630Δ*erm* wild-type (WT) and P*_tet_*-*dccA* strains, in which the diguanylate cyclase-encoding gene *dccA* is expressed from a chromosomally integrated anhydrotetracycline (ATc)-inducible promoter. Cultures were grown in TY medium with 25 or 250 ng/mL ATc, and c-di-GMP levels were quantified by LC-MS/MS. Bars represent means ± SD (*n* = 3 independent experiments). \*\*\*\**P* ≤ 0.0001 by a two-way analysis of variance (ANOVA) followed by a Sidak’s multiple comparisons test. Volcano plot of differentially expressed genes in the P*_tet_*-*dccA* strain compared to the WT strain. RNAseq was performed from early stationary phase cultures grown in TY with 250 ng/mL (*n* = 4 independent total RNA preparations). Selected genes are indicated. **(C)** Number of sporulation-related genes repressed under high c-di-GMP conditions, grouped according to their dependence on Spo0A, σ^E^, σ^F^, σ^G^, or σ^K^. The Spo0A and sporulation-associated sigma factors regulons are based on those reported in ^74^. Bars show the proportion of each regulon affected.

To assess the global impact of c-di-GMP, we compared the transcriptomes of wild type 630Δ*erm* and P*_tet_*-*dccA* strains grown to early stationary phase in TY with ATc (Supplementary Data 1). Using thresholds of a log_2_ fold change of greater than 2 or less than −2 and *P_adj_* ≤ 0.05, we identified 239 genes or RNA motifs differentially expressed under increased c-di-GMP conditions. Among them, 52 were upregulated and 187 were downregulated (Fig. 1B and Supplementary Table 1). As expected, *dccA* was the most strongly induced gene under elevated c-di-GMP conditions. Interestingly, the adjacent *CD1419* gene, which encodes another diguanylate cyclase^13^, was also strongly upregulated.

However, RNA-seq read coverage revealed a clear drop in signal between the two coding sequences (Supplementary Fig. 1), arguing against transcriptional readthrough. In addition, previous genome-wide transcription start site mapping identified an independent promoter upstream of *CD1419* ^43^, suggesting that *CD1419* is independently transcribed but highly responsive to increased c-di-GMP levels.

Other strongly induced genes (>10-fold increase) included *pilA1* (encoding the major pilin of type IV pili), *CD2831* (a cell-surface protein), and *CD1990* (an SH3 domain protein), all preceded by c-di-GMP-responsive riboswitches and known to be positively regulated by c-di-GMP ^14,18^. Expression of the predicted c-di-GMP phosphodiesterase *CD1616* ^13^ was also induced (4.2-fold), likely reflecting feedback regulation. Conversely, the most strongly repressed genes were *CD1424, CD1990.3, CD2309,* and *CD3368.2*, exhibiting 21- to 52-fold decreases in transcript abundance. These genes encode conserved hypothetical proteins and are preceded by class I c-di-GMP riboswitches (Cdi1_8, Cdi1_9, Cdi1_10, and Cdi1_11) ^14,18^, whose transcript levels also declined. The class I riboswitch Cdi1_6, not linked to an annotated CDS, was similarly repressed.

Strikingly, 81 sporulation-related genes, dependent on Spo0A and/or sporulation sigma factors σ^F^, σ^E^, σ^G^, and σ^K^, were downregulated under high c-di-GMP conditions (Supplementary Table 2). They included early sporulation genes such as the forespore-specific *spoIIAA-spoIIAB-sigF* operon and *spoIIE* (encoding the σ^F^-activating phosphatase), as well as the mother cell-specific gene *sigE* ^40^. In total, 29% of Spo0A- and 21% of σ^E^-dependent transcripts were repressed (Fig. 1C), consistent with previous evidence that c-di-GMP inhibits early sporulation in *C. difficile* ^33^.

### Class I riboswitches precede genes encoding a family of nearly identical small proteins

Transcriptome analyses revealed that c-di-GMP strongly affected several class I riboswitches (Cdi1_6, Cdi1_8, Cdi1_9, Cdi1_10, and Cdi1_11) that were not previously characterized.

Four lie upstream of conserved hypothetical protein genes, while Cdi1_6 lacked an annotated associated coding sequence. Bioinformatic analyses, however, identified a downstream open reading frame (ORF) encoding a 58-amino acid (AA) protein with a predicted ribosome-binding site 11 nucleotides upstream of the start codon (Fig. 2A). To test whether this ORF encodes a protein, we expressed it with an *in-frame* human influenza hemagglutinin (HA) tag under the ATc-inducible P*_tet_* promoter (p/CD1981-HA). Immunoblotting with anti-HA antibodies detected an ∼8 kDa product in cells grown with the inducer ATc but not in the control strain harboring an empty plasmid (Fig. 2B). These data confirm translation of a genuine small protein, designated CD1980.2.

**Fig. 2.**
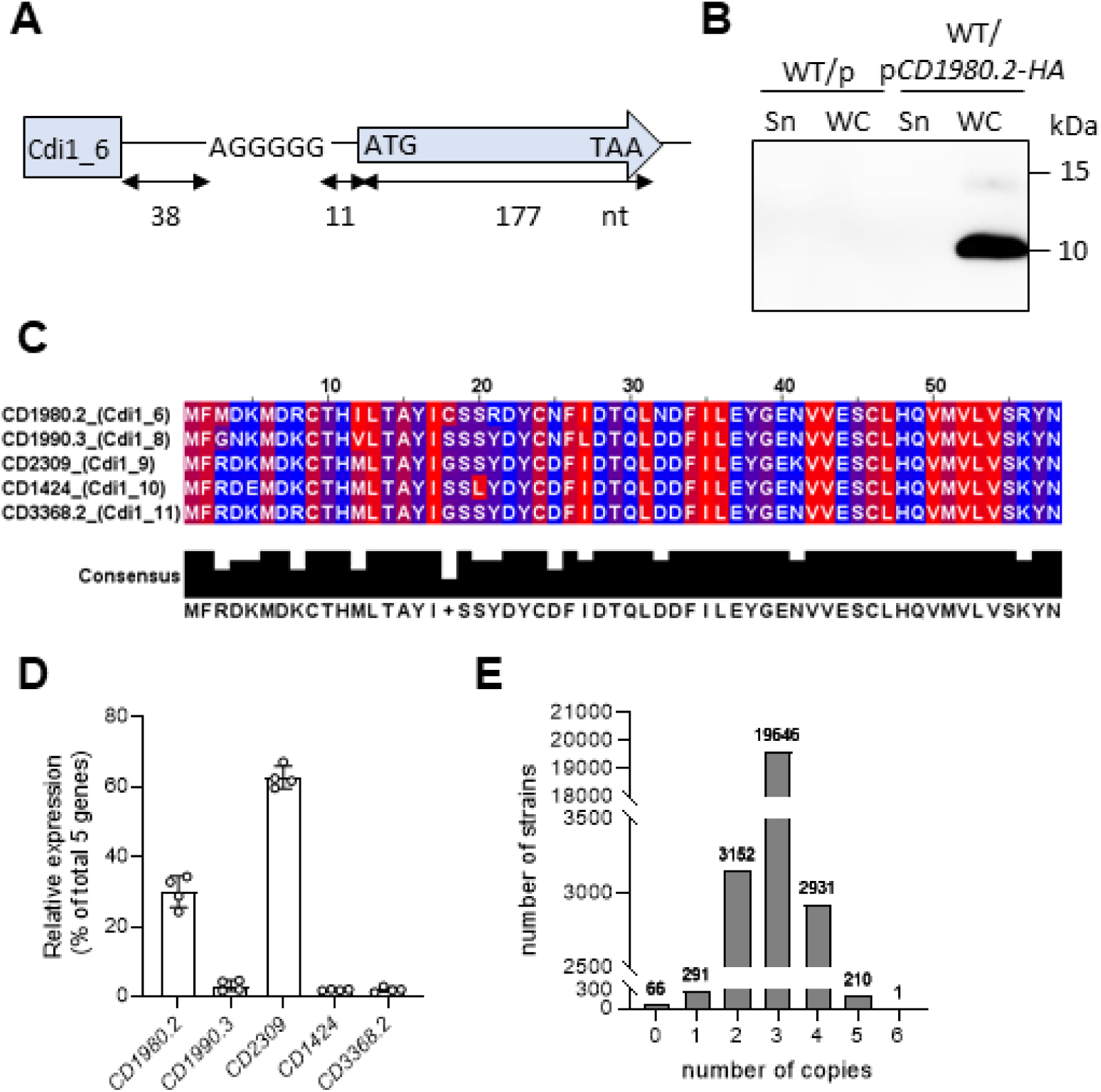
Class I c-di-GMP riboswitches control expression of a conserved family of small proteins in *C. difficile*. **(A)** Schematic representation of the Cdi1_6 riboswitch locus. The downstream CDS (*CD1980.2*) is depicted as an arrow with the start (ATG) and the stop (TAA) codons, and the size of the CDS is indicated. The putative RBS sequence (AGGGGG) is represented and the number of nucleotides between the RBS and the ribowitch and between the RBS and the CDS are indicated. **(B)** Detection of CD1980.2_HA_ by immunoblotting. Supernatant (Sn) and whole cell extract (WC) of the wild-type (WT) strain carrying either an empty plasmid (p) or a plasmid expressing CD1980.2_HA_ from the P*_tet_* promoter (p*CD1980.2-HA*) were analyzed by Western blotting with anti-HA antibodies. Strains were cultivated in the presence of 250 ng/mL ATc. Data are representative of three independent experiments. **(B)** Alignment of the five small proteins. The name of the c-di-GMP-responding riboswitch preceding each protein-encoding gene is indicated. The hydrophobic and positively charged amino acids are indicated in red and blue, respectively. **(D)** Estimation of the relative mRNA abundance of the five small protein-encoding genes based on RNA-seq reads from the WT strain grown in TY with 250 ng/mL ATc. Bars represent means ± SD (*n* = 4 independent experiments) **(E)** Determination of the number of small protein-encoding genes over 26,000 strains of *C. difficile*.

CD1980.2 lacks recognizable domains or non-*C. difficile* homologs. However, four highly similar 58 AA paralogs (76-88% identity) were identified (Fig. 2C), each located downstream of riboswitches Cdi1_8 to Cdi1_11, revealing a family of five nearly identical riboswitch-associated small proteins. Although these riboswitch-associated regions were previously reported to respond to elevated c-di-GMP levels ^18^, three downstream genes were annotated only as hypothetical proteins and no coding sequences were assigned downstream of Cdi1_6 and Cdi1_10. Our analyses show that all five riboswitches are associated with genes encoding conserved 58-aa membrane-associated proteins. RNA-seq read mapping indicated uneven expression with *CD2309* contributing ∼63% of the total reads assigned to these five loci, *CD1980.2* ∼30%, and the remaining three copies each <5%. (Fig. 2D). A truncated homolog (35 AA) associated with Cdi1_7 was also detected, disrupted by insertion of CRISPR 12 (Supplementary Fig. 2). Comparative genomics across >26,000 *C. difficile* genomes using the Diamond BLASTX function against the AllTheBacteria database (https://allthebacteria.readthedocs.io/en/latest/; v2) showed at least two intact copies in >98% of strains (≥90% identity; ≥50 AA) (Fig. 2E), highlighting the strong conservation and multicopy nature of this small-protein family.

### Additional CD1980.2 paralogs are riboswitch-independent

Beyond the five riboswitch-associated paralogs, three additional CD1980.2 paralogs were identified in *C. difficile* 630Δ*erm*: CD1667.1 (58 AA; 76% identity) and two 60-AA proteins, CD1511.1 and CD2386.1 (50-55% identity) (Supplementary Fig. 3A-C). None is preceded by a predicted c-di-GMP riboswitch. Nevertheless, RNA-seq revealed that *CD1511.1* and *CD2386.1* transcripts decreased 3.7- and 7-fold, respectively, under high c-di-GMP conditions (Supplementary Table 1), indicating riboswitch-independent regulation by c-di-GMP. By contrast, *CD1667.1* was barely transcribed under either condition (Supplementary Fig. 3D). Previous work suggested that *CD1667.1* expression is silenced by the *cis*-encoded antisense RNA CD630_n00620, renamed SpoZ ^14,44^. Consistent with this model and RNA chaperone Hfq RIP-seq profile ^45^, abundant reads mapping to *spoZ* overlapped the *CD1667.1* ORF in both conditions of our RNA-seq dataset (Supplementary Fig. 3D).

Expression of *spoZ* and *CD1511.1* is activated by the RgaR-RgaS two-component system through the binding of RgaR to the VirR-like binding site in the promoter region ^44^. A putative VirR-binding site was also identified upstream of *CD2386.1* (Supplementary Fig. 3E), suggesting that all three riboswitch-independent paralogs are controlled directly or indirectly by RgaR. We therefore examined whether overexpression of the diguanylate cyclase DccA affected the transcription of the other reported RgaR-regulated genes. The *agrB1/D1* operon, *CD2098* and *CD0587* displayed modest decreases in expression (2.2 to 4.3-fold), whereas *spoZ* expression was not significantly affected (Supplementary Table 1).

### The family of small proteins is membrane-associated and c-di-GMP-regulated

Sequence analysis revealed a C-terminal hydrophobic domain in the small protein family (Fig. 2C and Supplementary Fig. 3A-C), suggesting membrane association. Structural comparison of AlphaFold3-predicted models of the five paralogs (CD2309, CD1424, CD1980.2, CD1990.3, and CD3368.2) revealed a conserved four-helix bundle (global RMSD < 0.5 Å; Supplementary Fig. 4A). Helices 1 and 3 form an antiparallel core (interhelical angles 143.4–159.3°), while helices 2 and 4 adopt more oblique orientations (Supplementary Fig. 4B).

Independent alignment of the C-terminal Helix 4 (residues 40–57) showed striking structural conservation (mean RMSD = 0.21 Å). Furthermore, molecular lipophilicity potential (MLP) mapping revealed a focal hydrophobic patch (MLP_max_ > 22) restricted to the outward-facing surface of Helix 4 (Supplementary Fig. 4C), consistent with a role in membrane association or insertion. Coulombic electrostatic analysis further revealed a concentrated electropositive patch on the exposed helical core (Helices 1–3), suggesting a potential interface for interaction with polyanionic ligands or acidic protein partners (Supplementary Fig. 4D). The Helix 4 consensus (residues 40–57) displays an *i*, *i*+4 Val/Cys/Met-enriched hydrophobic periodicity (V42–C46–V50–V54), suggestive of a transmembrane dimerization interface, and conservation of Cys46, raising the possibility of covalent stabilization (ipTM = 0.14).

Immunoblotting with anti-HA antibodies of subcellular fractions of *C. difficile* expressing CD1980.2-HA confirmed membrane localization (Fig. 3A). Because plasmid-based overexpression can artificially drive protein aggregation, we validated membrane localization at native expression levels. For this, we integrated a sequential peptide affinity (SPA) tag (3×FLAG and calmodulin-binding peptide) into the chromosome, *in-frame* upstream of the stop codon of *CD2309*. This locus was chosen as a prototype since *CD2309* transcripts were the most abundant among the paralogs (Fig. 2D). Immunoblotting confirmed that CD2309-SPA was enriched in the membrane fraction (Fig. 3B).

**Fig. 3.**
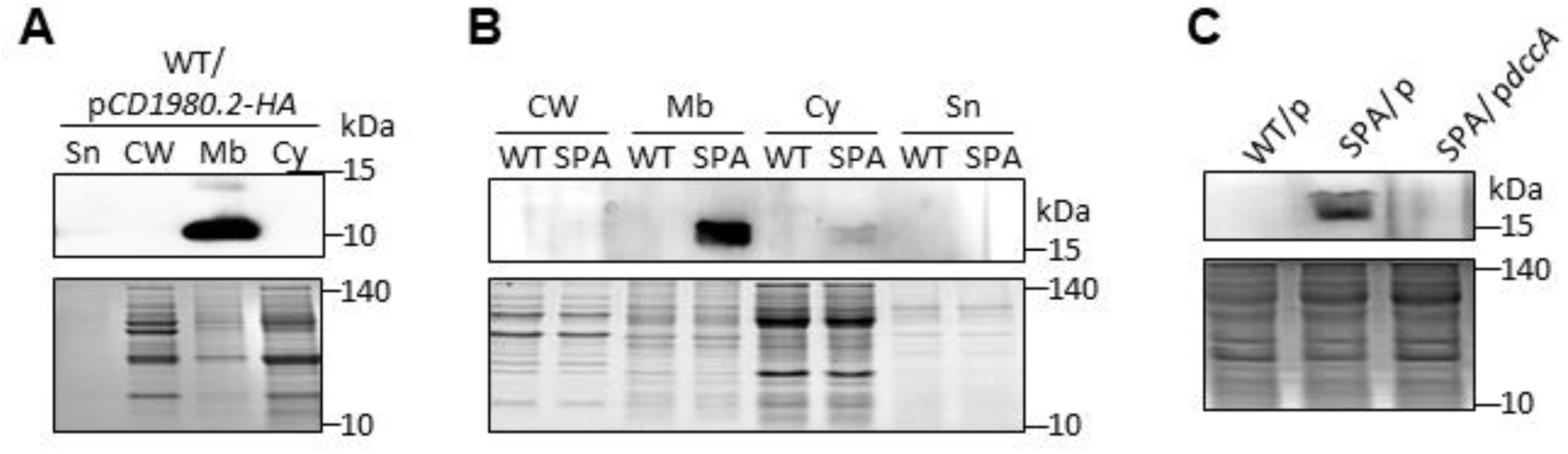
The small protein family is membrane-associated and negatively regulated by the c-di-GMP accumulation. **(A)** Supernatant (Sn), cell wall (CW), membrane (Mb), and cytosolic (Cy) compartments of *C. difficile* carrying a plasmid expressing *CD1980.2_HA_* from the P*_tet_* promoter (p*CD1980.2-HA*) were analyzed by Western blotting with anti-HA antibodies. Strains were cultivated in the presence of 250 ng/mL ATc. An InstantBlue-stained portion of the SDS-PAGE gel is shown as a loading control (lower panel). Data are representative of three independent experiments. **(B)** Subcellular fractions of a strain in which the SPA tag-encoding sequence was fused to *CD2309* into the chromosome (SPA) and of the isogenic wild-type (WT) strain were analyzed by Western blotting with anti-FLAG M2 antibodies. Strains were cultivated in the presence of 100 ng/mL ATc. An InstantBlue-stained portion of the SDS-PAGE gel is shown as a loading control (lower panel). Data are representative of three independent experiments. **(C)** Whole cell extracts of the WT strain carrying an empty plasmid (p) and of the SPA strain carrying either an empty plasmid or a plasmid expressing *dccA* from the P*_tet_* promoter (p*dccA*) were analyzed by Western blotting with anti-FLAG M2 antibodies. Strains were cultivated in the presence of 100 ng/mL ATc. An InstantBlue-stained portion of the SDS-PAGE gel is shown as a loading control (lower panel). Data are representative of three independent experiments.

To test c-di-GMP-dependent regulation, CD2309-SPA strains carrying either an empty vector or P*_tet_*-*dccA* were induced with ATc. CD2309-SPA abundance decreased markedly upon *dccA* overexpression (Fig. 3C), consistent with transcriptomic data. Together, these findings demonstrate that CD2309, and likely the entire family of riboswitch-associated small proteins, are membrane-associated and negatively regulated by c-di-GMP.

### C-di-GMP controls the expression of the small protein-encoding genes through interaction with their cognate riboswitches

To determine whether c-di-GMP directly regulates small protein expression through associated riboswitches, we constructed transcriptional and translational reporter fusions on plasmids. For *CD2309*, the promoter region, with or without the 5′ untranslated region (5′ UTR) harboring the Cdi1_9 riboswitch, was placed upstream of the *phoZ* reporter gene (Fig. 4A). The construct containing only the promoter displayed similar alkaline phosphatase activity in wild-type and P*_tet_*-*dccA* strains grown with 25 or 250 ng/mL Atc (Fig. 4B and Supplementary Fig. 5A), indicating that promoter activity is not affected by c-di-GMP. Inclusion of the 5′ UTR encompassing the Cdi1_9 riboswitch led to an unchanged activity in the wild type but a ∼2-fold increase in reporter activity under high c-di-GMP conditions (Fig. 4B and Supplementary Fig. 5A). A translational fusion produced a similar increase in activity under the same conditions (Fig. 4B and Supplementary Fig. 5A). A comparable trend was also observed with a translational fusion (Fig. 4B and Supplementary Fig. 5A) and with the Cdi1_6 riboswitch region of *CD1980.2* (Supplementary Fig. 5B), demonstrating that the riboswitch is required for c-di-GMP-dependent modulation.

**Fig. 4.**
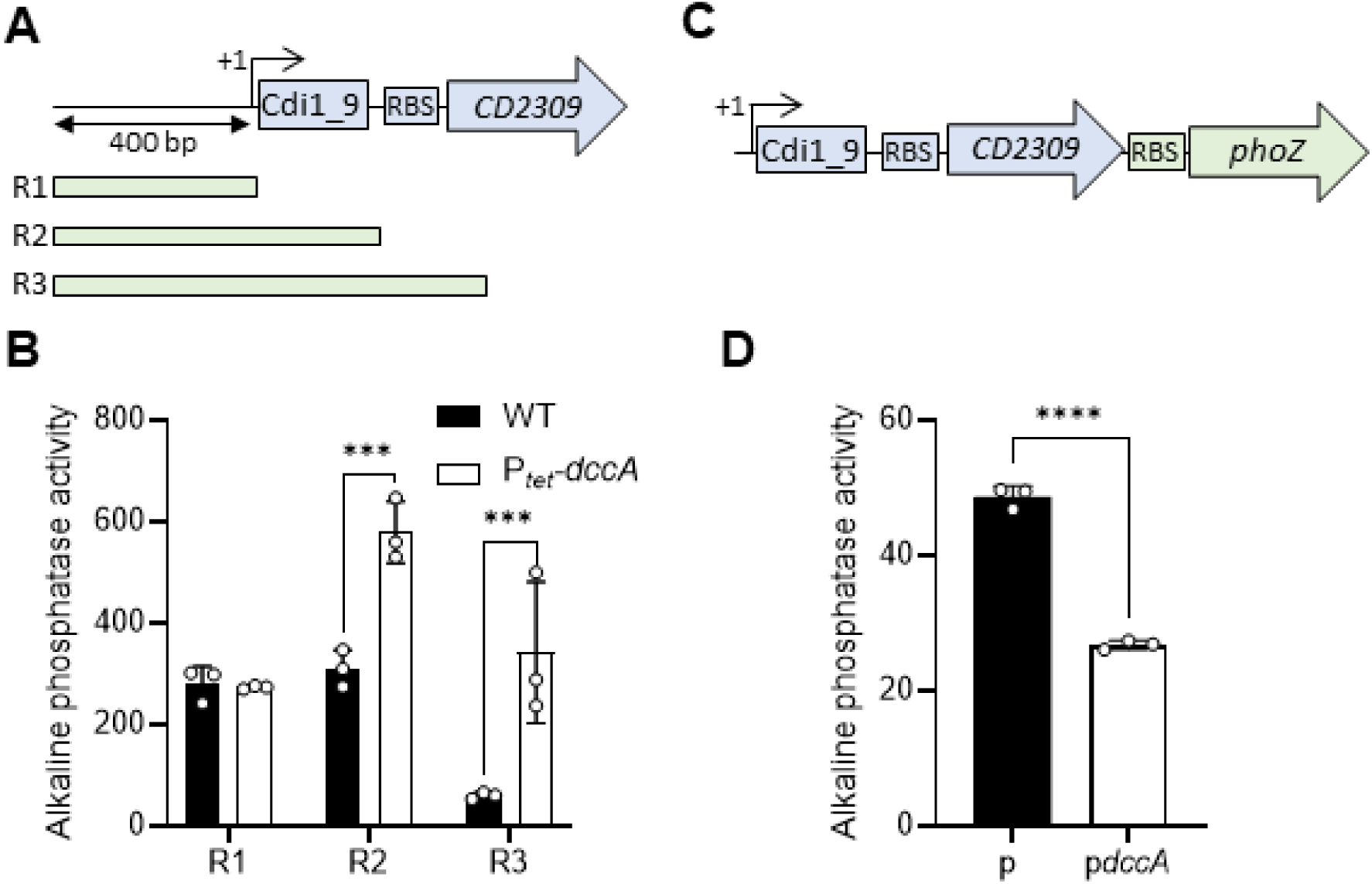
c-di-GMP controls CD2309 expression through direct regulation by the upstream c-di-GMP riboswitch. **(A)** Schematic of the Cdi1_9-CD2309 genomic region and of the location and size of the regions (R1 to R3) fused to the reporter gene *phoZ* on a plasmid. **(B)** Alkaline phosphatase activity of the Cdi1_9::*phoZ* reporter fusions of various lengths in the wild-type (WT) and the P*_tet_-dccA* strains, in which the diguanylate cyclase-encoding gene *dccA* is expressed from a chromosomally integrated anhydrotetracycline (ATc)-inducible promoter. Strains were grown in the exponential phase in TY with 250 ng/mL ATc. Bars represent means ± SD (*n* = 3 independent experiments). \*\*\**P* ≤ 0.001 by a two-way ANOVA followed by a Sidak’s multiple comparisons test. **(C)** Schematic of the *CD2309*::*phoZ* transcriptional fusion carried on the chromosome. The construct corresponds to the insertion of *phoZ* preceded by a consensus RBS directly downstream the *CD2309* CDS, which allows the translation of the reporter without being affected by the structure of the riboswitch. **(D)** Alkaline phosphatase activity of the *CD2309*::*phoZ* reporter fusion in the WT strain carrying either an empty plasmid or a plasmid expressing *dccA* from the P*_tet_* promoter (p*dccA*). Strains were grown in the exponential phase in TY with 250 ng/mL ATc. Bars represent means ± SD (*n* = 3 independent experiments). \*\*\*\**P* ≤ 0.0001 by an unpaired Student’s *t* test.

Notably, this increased reporter activity contrasts with our RNA-seq and immunoblot data, which showed reduced *CD2309* transcript and protein under high c-di-GMP conditions. This difference appears specific to the Cdi1_6 and Cdi1_9 riboswitches, as a control fusion containing the *flgB* promoter and its Cdi1_3 riboswitch, previously characterized as an “OFF” switch ^17^, displayed the expected repression under high c-di-GMP (Supplementary Fig. 5C). Rather than reflecting a technical artifact, these observations suggest that full regulatory activity of the Cdi1_6 and Cdi1_9 riboswitches requires the native chromosomal context, potentially relying on local genomic architecture or RNA-protein interactions absent from plasmid-based constructs. Consistent with this view, a chromosomal *CD2309*::*phoZ* fusion reproduced the expected repression under high c-di-GMP (Fig. 4C and 4D).

Taken together, these results support a model in which c-di-GMP interacts with class I riboswitches to trigger premature transcription termination of small protein-encoding mRNAs. Importantly, this regulation requires the native chromosomal context, highlighting the possible contribution of additional cis-acting elements to c-di-GMP-dependent control.

### Overexpression of CD1980.2 promotes sporulation

To explore the physiological role of this small protein family, the Cdi1_6*-CD1980.2* locus was placed under the control of the P*_tet_* promoter on a plasmid and introduced into the ΔCdi1_6-*CD1980.2* mutant strain. Transcriptomic changes were then assessed by RNA-seq, comparing this *CD1980.2* overexpressing strain to the ΔCdi1_6-*CD1980.2* mutant carrying an empty plasmid (Supplementary Data 2). Cultures were grown to the late exponential phase in TY medium in the presence of the inducer ATc. Pairwise transcriptome comparisons revealed 124 genes differentially expressed (log_2_ fold change >2 or < -2; adjusted *P_adj_* < 0.05) upon *CD1980.2* overexpression. Among them, 121 were upregulated and only 3 were downregulated (Fig. 5A and Supplementary Table 3). Notably, 83 of the induced genes were associated with sporulation, many displaying substantial expression changes (Supplementary Table 4). For instance, the spore protein-encoding gene *sspB* showed up to a 261-fold increase. This transcriptional activation occurred despite culture conditions that are generally unfavorable for sporulation induction. Closer analysis of the induced genes highlighted a striking enrichment of transcripts dependent on the late-acting sigma factors σ^G^ and σ^K^ (63% and 74% of their regulons, respectively), whereas transcripts dependent on Spo0A, σ^E^, and σ^F^ were less represented (10, 9, and 13%, respectively) (Fig. 5B). These results indicate that *CD1980.2* overexpression promotes progression well beyond the initiation of sporulation, driving the cells toward the late stages of spore formation.

**Fig. 5.**
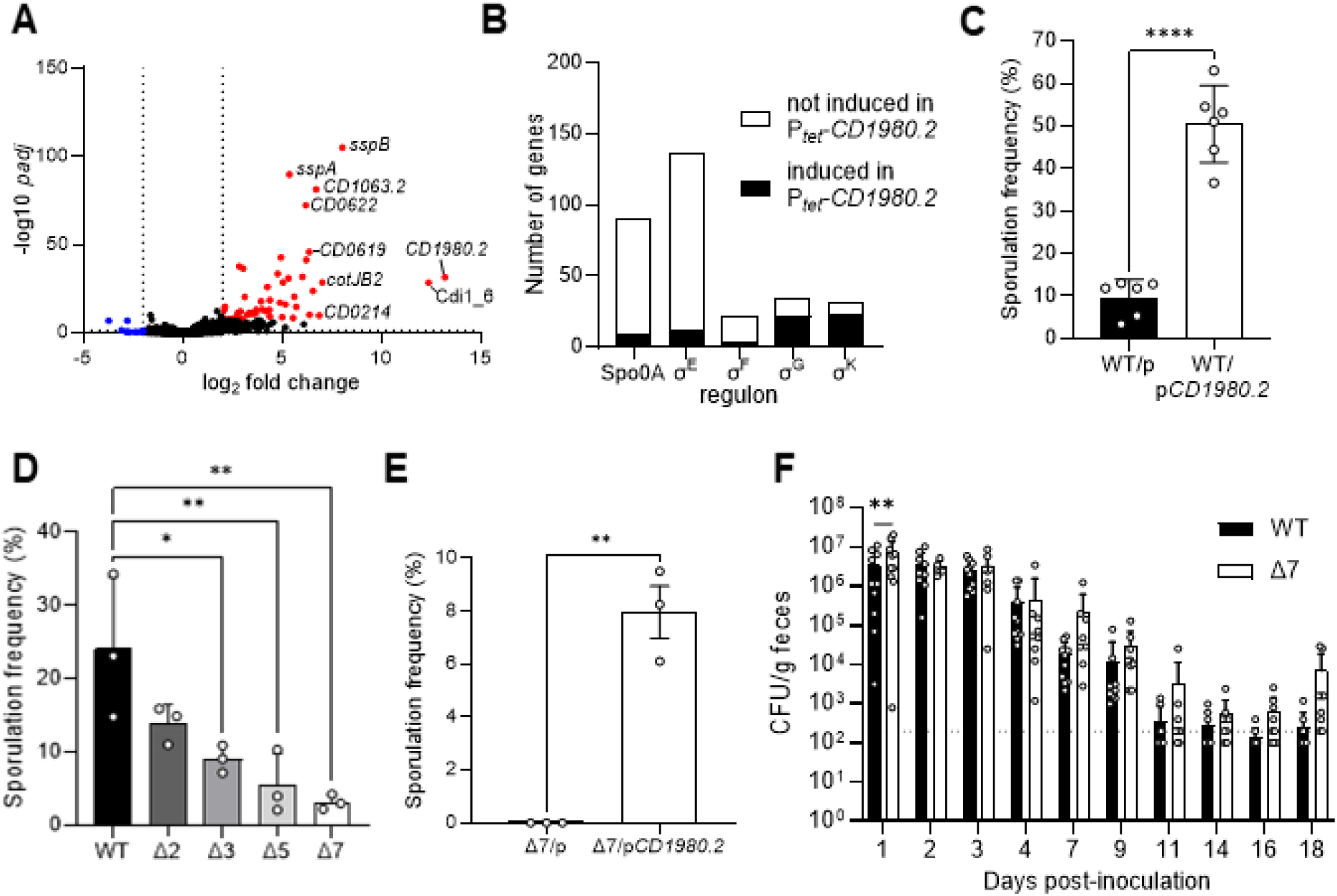
A conserved family of c-di-GMP-regulated small proteins drives sporulation in *C. difficile*, with cumulative effects on spore formation but no impact on colonization. (A) Volcano plot of differentially expressed genes in the ΔCdi1_6-*CD1980.2* strain carrying a plasmid expressing Cdi1_6*-CD1980.2* under the control of the P*_tet_* promoter compared to the same strain carrying an empty plasmid. RNAseq was performed from early stationary phase cultures grown in TY with 250 ng/mL ATc (*n* = 4 independent total RNA preparations). Selected genes are indicated. **(B)** Number of sporulation-related genes induced under Cdi1_6*-CD1980.2* (P*_tet_*-*CD1980.2*) expression conditions, grouped according to their dependence on Spo0A, σ^E^, σ^F^, σ^G^, or σ^K^. The Spo0A and sporulation-associated sigma factors regulons are based on those reported in ^74^. Bars show the proportion of each regulon affected. **(C)** Ethanol-resistant spore formation of the WT carrying an empty plasmid (p) or a plasmid expressing Cdi1_6*-CD1980.2* under the control of the P*_tet_* promoter (p*CD1980.2*), grown for 24h on 70:30 agar in the presence of 100 ng/mL ATc. Bars represent means ± SD (*n* = 6 independent experiments). \*\*\*\**P* ≤ 0.0001 by an unpaired Student’s *t* test. (D) Ethanol-resistant spore formation of a strain lacking two (Δ2), three (Δ3), five (Δ5) or seven (Δ7) small protein copies and of the isogenic WT strain grown for 24h on 70:30 agar. Bars represent means ± SD (*n* = 3 independent experiments). \**P* ≤ 0.05 and \*\**P* ≤ 0.01 by a one-way ANOVA followed by a Dunnett’s multiple comparisons test. **(E)** Ethanol-resistant spore formation of the Δ7 mutant strain carrying an empty plasmid (p) or a plasmid expressing Cdi1_6*-CD1980.2* under the control of the P*_tet_* promoter (p*CD1980.2*) grown for 24h in liquid 70:30 medium in the presence of 250 ng/mL ATc. Bars represent means ± SD (*n* = 3 independent experiments). \*\**P* ≤ 0.01 by an unpaired Student’s *t* test. (F) Antibiotic-treated female C57BL/6 mice were inoculated with 10^5^ spores of wild type 630Δ*erm* (WT) or Δ7. Feces were collected on the indicated days and plated on selective medium to assess the total number of CFUs. Bars represent means ± SD (*n* = 10 mice). \*\**P* < 0.01 by a two-way ANOVA followed by a Sidak’s multiple comparisons test.

We next compared sporulation frequencies between the strains 630Δ*erm* carrying the plasmid expressing Cdi1_6*-CD1980.2* or harboring an empty plasmid. Both strains were grown for 24 hours on 70:30 sporulation agar supplemented with ATc, and the ratio of spores to vegetative cells was determined. Overexpression of *CD1980.2* led to an approximately 5-fold increase in spore formation relative to the control strain (Fig. 5C). An even stronger hypersporulation phenotype was observed in liquid 70:30 medium (Supplementary Fig. 6A), indicating that its sporulation-promoting activity is robust across different sporulation conditions. Thus, ectopic expression of *CD1980.2* is sufficient to promote sporulation in *C. difficile*, suggesting that this small protein functions as a positive regulator of the sporulation pathway. To assess whether this effect is conserved in a clinically relevant genetic background, we next tested CD1980.2 in the epidemic isolate UK1. Under the conditions tested, the UK1 strain carrying the empty vector already exhibited a high sporulation efficiency in liquid 70:30 medium (∼18%), comparable to that observed for the *CD1980.2*-overexpressing 630Δ*erm* strain (Supplementary Fig. 6A). Overexpression of *CD1980.2* in UK1 did not further increase sporulation. Thus, the effect of *CD1980.2* overexpression appears to be dependent on the strain background and/or the basal sporulation capacity of the strain.

Given that *CD1980.2* expression is directly repressed by c-di-GMP and that its ectopic expression is sufficient to promote sporulation in strain 630Δ*erm*, we next examined whether sporulation is accompanied by changes in c-di-GMP-dependent gene expression. Consistent with this model, qRT-PCR analyses under sporulation conditions revealed only minor changes in the expression of representative c-di-GMP-responsive genes ^15^ (Supplementary Fig. 6B-C). These data suggest that any variation in c-di-GMP signaling during sporulation is likely subtle and/or spatially localized.

### The small proteins are redundantly critical for sporulation but are dispensable for gut colonization

Given their high sequence similarity, we hypothesized that the small proteins act redundantly during sporulation. To test this, we generated a multiple-deletion strain lacking all five 58-AA protein-encoding genes as well as *CD1511.1* and *CD2386.1* (Δ7 mutant; Supplementary Fig. 7). When grown on 70:30 sporulation agar for 24h, the Δ7 mutant displayed a severe sporulation defect compared with the wild-type (Fig. 5D). Intermediate mutants lacking two (Cdi1_6 and Cdi1_9 loci), three (Cdi1_6, Cdi1_8 and Cdi1_9 loci) or five (all riboswitch-associated loci) small protein gene copies exhibited proportionally reduced sporulation efficiencies (Fig. 5D), demonstrating that the small proteins contribute cumulatively and redundantly to sporulation. Complementation of the Δ7 mutant with *CD1980.2* under the control of the P*tet* promoter restored sporulation and significantly increased sporulation efficiency compared with the vector control (Fig. 5E). These results confirm that the sporulation defect of the Δ7 strain is directly attributable to loss of CD1980.2 family proteins.

To better understand the limited impact of the double mutant, we next examined whether paralog expression profiles differed under sporulation-inducing conditions. Because CD1980.2 and CD2309 represented the two most abundant paralogs during growth in TY medium (Fig. 2D), we asked whether this distribution changed under sporulation conditions. RNA-seq of the wild-type strain grown for 16 h on 70:30 agar (Supplementary Data 3) showed that CD1980.2 and CD2309 remained the major transcripts (∼48% and ∼43%, respectively Supplementary Fig. 8A). qRT-PCR further showed no compensatory increase in expression of the remaining paralogs in representative deletion mutants (Supplementary Fig. 8B). Thus, the modest phenotype of the double mutant is unlikely to result from transcriptional compensation. Instead, low-abundance paralogs may suffice to support sporulation, or paralogs may contribute unequally to this process.

We next examined whether this *in vitro* sporulation defect affects intestinal colonization. In an antibiotic-treated conventional mouse model, animals were infected orally with 10^5^ spores of either the wild type or the Δ7 mutant, and fecal *C. difficile* CFUs were monitored over 18 days (Fig. 5F). Neither strain caused any detectable clinical impact in the animals, and both exhibited comparable colonization levels throughout the experiment, indicating that the small proteins are dispensable for intestinal colonization. These results indicate that the small proteins primarily influence sporulation and potentially spore-mediated transmission, rather than the establishment or maintenance of intestinal colonization.

### c-di-GMP represses sporulation through the control of the small protein expression

Previous work showed that elevated c-di-GMP concentrations inhibit *C. difficile* sporulation ^33,34^, but the mechanism remained unclear. We hypothesized that repression of the small proteins mediates this effect. To test this, we first compared the sporulation frequency of the P*_te_*_t_-*dccA* strain, which overproduces c-di-GMP, to the Δ7 multiple-deletion strain. On 70:30 agar with ATc, P*_te_*_t_-*dccA* displayed a strong reduction in spore formation, comparable to the Δ7 mutant (Fig. 6A).

**Fig. 6.**
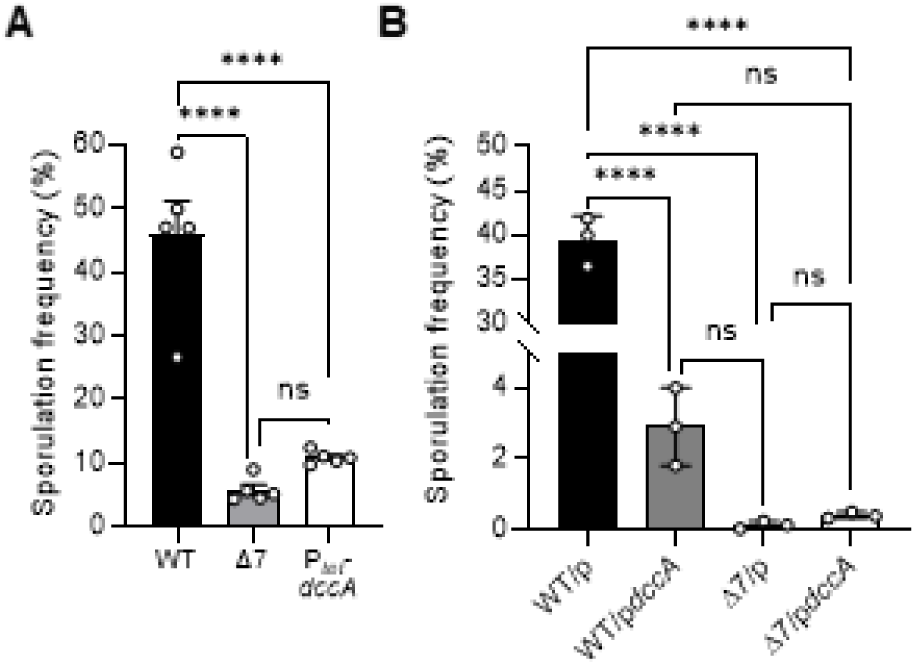
c-di-GMP inhibits sporulation through the small protein family in *C. difficile*. **(A)** Ethanol-resistant spore formation of a strain lacking the seven small protein copies (Δ7), of the P*_tet_*-*dccA* strain, in which the diguanylate cyclase-encoding gene *dccA* is expressed from a chromosomally integrated anhydrotetracycline (ATc)-inducible promoter and of the isogenic wild-type strain (WT) grown for 24h on 70:30 agar. Bars represent means ± SD (*n* = 5 independent experiments). \*\*\*\**P* ≤ 0.0001 and ns = not significant by a one-way ANOVA followed by a Tukey’s multiple comparisons test. (B) Ethanol-resistant spore formation of WT and Δ7 carrying either an empty plasmid (p) or a plasmid expressing *dccA* from the P*_tet_* promoter (p*dccA*) grown for 24h on 70:30 agar. Bars represent means ± SD (n = 3 independent experiments). \*\*\*\**P* ≤ 0.0001 and ns = not significant by a one-way ANOVA followed by a Tukey’s multiple comparisons test.

Next, we introduced the p*dccA* plasmid into both the wild-type and the Δ7 backgrounds and measured sporulation frequencies alongside appropriate empty-vector controls (Fig. 6B). As expected, either c-di-GMP overproduction or deletion of the small proteins significantly reduced sporulation, but the combination did not further decrease spore formation. We cannot exclude that the very low sporulation level in the Δ7 mutant limits detection of any residual c-di-GMP effect. Nevertheless, the progressive decrease in sporulation across intermediate deletion strains, together with overexpression and transcriptional data, strongly suggests that these small proteins are the main mediators of c-di-GMP-dependent inhibition. Specifically, our results support a model in which c-di-GMP represses sporulation initiation by reducing the expression of the small protein-encoding genes, linking intracellular second messenger levels to developmental control in *C. difficile*. To place these findings in the broader context of c-di-GMP signaling in *C. difficile*, we propose a schematic overview integrating the small protein-mediated control of sporulation with previously characterized regulatory outputs (Fig. 7).

**Fig. 7.**
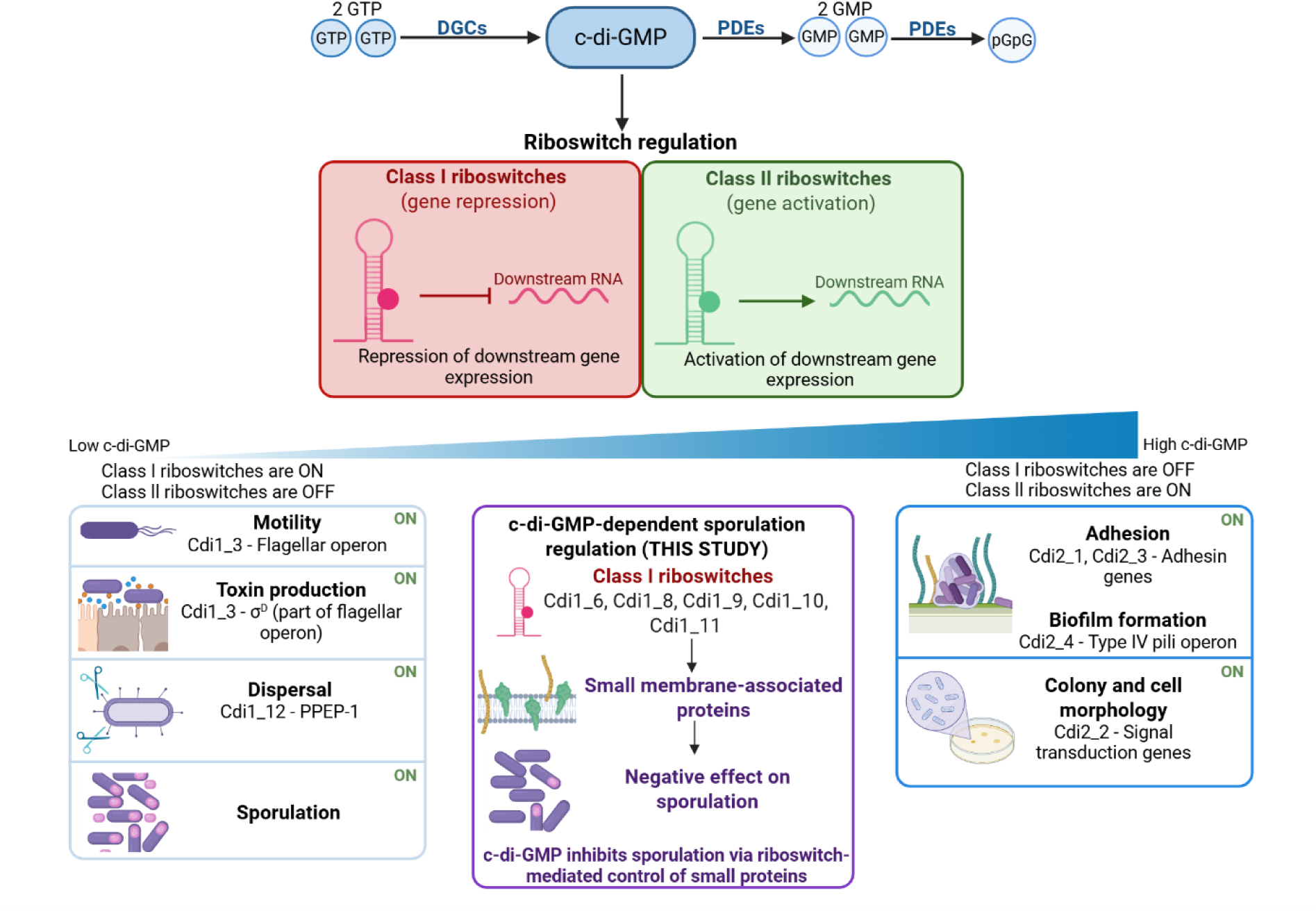
c-di-GMP signaling coordinates lifestyle transitions and inhibits sporulation via a riboswitch-dependent pathway in *Clostridioides difficile*. c-di-GMP is a central second messenger synthesized by diguanylate cyclases (DGCs) and degraded by phosphodiesterases (PDEs). c-di-GMP coordinates the transition between motile/virulent (low c-di-GMP) and sessile (high c-di-GMP) states in *C. difficile* through riboswitch-mediated gene regulation. Two major classes are present: class I riboswitches, which typically repress downstream gene expression upon c-di-GMP binding, and class II riboswitches, which activate gene expression ^16,17^. While low c-di-GMP promotes motility, dispersal mechanisms, toxin production, and sporulation, high c-di-GMP represses these processes and induces adhesion, biofilm formation, and bacterial elongation ^15^. This study reveals a novel pathway in which c-di-GMP inhibits sporulation via class I riboswitch-dependent regulation of small membrane-associated proteins, providing a mechanistic link between c-di-GMP signaling and developmental control.

## DISCUSSION

Cyclic di-GMP is a central bacterial second messenger that coordinates a wide range of physiological processes, including motility, biofilm formation, virulence, and developmental pathways ^1,2,46^. In *C. difficile*, previous studies established that elevated c-di-GMP concentration inhibits sporulation ^33,34^, but the molecular effectors linking this second messenger to the developmental machinery were unknown. Here, we identify a conserved family of small, membrane-associated proteins as critical intermediates connecting c-di-GMP signaling to sporulation. Phenotypic analyses demonstrate that they promote spore formation in a cumulative and redundant manner. Functionally redundant paralogs can enhance transcript abundance, partition ancestral functions, or acquire novel roles ^47,48^. In the absence of selective pressure, however, most duplicates are rapidly lost, and bacteria retain only those duplications essential for physiology or adaptation ^47,49^. The conservation of multiple copies of these small protein genes across *C. difficile* lineages, including epidemic strains, strongly argues for their importance in a fundamental physiological process.

Our data are consistent with a model in which c-di-GMP represses sporulation, at least in part, by downregulating the expression of these small proteins. However, additional genetic analyses will be needed to fully resolve the organization of this regulatory pathway. The available evidence further suggests that this regulation occurs primarily at the transcriptional level via the associated riboswitches, potentially through premature termination, a process that functions optimally on the chromosome rather than on multicopy plasmids. Importantly, our results do not indicate that a decrease in c-di-GMP is required to trigger expression of these genes, as they are already expressed during exponential growth. Instead, these small proteins likely act as permissive factors for sporulation, whose presence is required prior to entry into the developmental program. In this context, c-di-GMP would primarily function as a negative regulator, preventing sporulation by repressing the expression of these proteins when its intracellular concentration is elevated. Consistent with this model, we observed only modest changes in the expression of representative c-di-GMP riboswitch-associated genes during growth under sporulation conditions (Supplementary Fig. 6B). These observations suggest that c-di-GMP variations during sporulation are likely subtle and spatially restricted, consistent with models of localized c-di-GMP signaling ^50^. Notably, three homologs of this family, namely CD1667.1, CD1511.1, and CD2386.1, lack detectable riboswitches, suggesting alternative mechanisms of regulation. Among them, CD1667.1 has previously been shown to be negatively regulated by the antisense sRNA SpoZ ^44^. Knockdown of *spoZ* increased spore formation, whereas overexpression of CD1667.1 reduced sporulation. This counterintuitive relationship may reflect SpoZ acting on multiple targets, complicating the interpretation. Interestingly, the effect of CD1667.1 contrasts with that of CD1980.2. While CD1667.1 reduces sporulation, CD1980.2 promotes hypersporulation despite the two proteins sharing 76% identity. This observation raises the possibility that some paralogs exert opposing functions or compete for common regulatory pathways.

The two-component system RgaS/RgaR provides an additional regulatory layer. RgaR directly binds a consensus site upstream of *spoZ*, activating its transcription ^44^. Accordingly, Δ*rgaS-R* mutants fail to express *spoZ*, resulting in unchecked CD1667.1 accumulation and decreased sporulation. RgaR binding sites were also detected upstream of four other genes, including *CD1511.1* ^44^. In agreement, *CD1511.1* expression decreased in a Δ*rgaR* mutant, yet its knockdown did not alter sporulation frequency, consistent with our finding that multiple paralogs must be deleted to reveal a phenotype. Strikingly, we also identified a near perfect RgaR binding site upstream of *CD2386.1*, showing that the three homologs of this family lacking detectable riboswitches are likely controlled directly or indirectly by RgaR. Although no riboswitches are predicted upstream of *CD1511.1* and *CD2386.1*, our data nevertheless indicate that their expression responds to c-di-GMP levels. In addition, our RNA-seq analysis revealed that c-di-GMP accumulation resulted in modest decreases in the expression of several other reported RgaR targets, including the agr*D1*, *CD2098* and *CD0587*, whereas spoZ was not affected (Table S1). While the underlying mechanism remains unresolved, these moderate effects suggest that elevated c-di-GMP levels may influence parts of the RgaR regulon, either through a direct effect on RgaR activity or through indirect regulatory pathways.

Small proteins are increasingly recognized as important regulators in bacterial physiology. Although historically overlooked due to annotation challenges and detection limitations, small proteins, typically defined as polypeptides of fewer than 50-100 amino acids, can interact with nucleic acids, larger proteins, and membranes to control diverse processes, including stress responses, cell division, transport, and sporulation ^51–55^ ^56,57^. In several model organisms, small proteins directly modulate sporulation. For example, in *B. subtilis*, the small protein Sda inhibits the sporulation kinase KinA, acting as a checkpoint to prevent initiation under conditions of DNA stress ^58,59^, while the small protein MciZ blocks FtsZ polymerization to control mother cell division during spore formation ^60,61^. Similarly, small proteins such as SpoVM and SpoIVA regulate spore coat assembly, ensuring proper forespore encapsulation and resistance to environmental stresses ^62–64^. In *C. difficile*, the family of small membrane-associated proteins identified here adds a new layer to this regulatory landscape, acting redundantly to promote sporulation under appropriate physiological conditions. Our study establishes a robust link between c-di-GMP, small proteins, and sporulation, yet the molecular basis of this regulation remains unclear. Whether these small proteins directly interact with sporulation regulators, membrane components, or other signaling elements is not yet known. Direct experimental validation using truncated variants of the small proteins would be valuable to assess the role of the C-terminal helix in membrane association and protein function. Although small, hydrophobic proteins are notoriously difficult to detect and characterize ^65,66^, further biochemical and structural studies will be needed to identify their molecular partners and clarify how c-di-GMP-responsive riboswitches integrate into *C. difficile*’s developmental network.

In summary, our work identifies a conserved family of c-di-GMP-regulated small proteins as key effectors of sporulation in *C. difficile*. Their redundancy, integration into both riboswitch-and RgaR-mediated regulatory networks, and strong impact on sporulation efficiency highlight their importance for the developmental biology of this pathogen. Given the role of spores in persistence and recurrence, deciphering the molecular mechanisms of these small proteins could open new avenues for therapeutic strategies aimed at controlling *C. difficile* infection.

## METHODS

### Bacterial strains and growth conditions

The bacterial strains used in this study are listed in Supplementary Table 5. *E. coli* strains were grown aerobically at 37°C in Luria Bertani (LB) broth (Lennox, Sigma) supplemented with ampicillin (100 µg/mL) or chloramphenicol (15 µg/mL), when necessary. *C. difficile* strains were cultured anaerobically in an anaerobic workstation (Jacomex) under an atmosphere containing 5% H₂, 5% CO₂, and 90% N₂. *C. difficile* strains were routinely cultured in TY, or Brain Heart Infusion media (BHI, BD Difco). When necessary, cefoxitin (8 µg/mL), cycloserine (250 µg/mL) and thiamphenicol (7.5 µg/mL) were added to *C. difficile* cultures. The non-antibiotic analog anhydrotetracycline (ATc) (Sigma-Aldrich) was used to induce the *P_tet_* promoter in pRPF185 vector derivatives ^67^.

### Plasmids and strains construction

All plasmids and oligonucleotides utilized in this study are listed in Supplementary Tables 5 and 6, respectively.

A toxin-mediated allele exchange method was used for gene deletion in *C. difficile* ^32^. Briefly, allele exchange cassettes were designed with ∼800 bp homology arms flanking the target deletion region. These homology arms were amplified by PCR using genomic DNA from *C. difficile* strain 630Δ*erm* and subsequently cloned into the PmeI site of the pseudo-suicide allele-coupled exchange (ACE) vector pMSR via NEBuilder HiFi DNA Assembly (New England Biolabs). All plasmids derived from pMSR were first transformed into *E. coli* NEB10β, and inserts were confirmed by sequencing. The plasmids were then introduced into *E. coli* HB101(RP4) and transferred into *C. difficile* strains by conjugation. Transconjugants were selected on BHI medium supplemented with cycloserine, cefoxitin, and thiamphenicol. Toxin-mediated allelic exchange was carried out as described previously ^32^, and all strains were validated by locus-specific amplification. The deletions and the absence of unintended mutations in the Δ7 mutant were confirmed by whole-genome sequencing.

To insert the SPA tag sequence or *phoZ* into the chromosome, homology arms of ∼800 bp were designed, flanking the region where insertion was desired. These homology arms were PCR amplified using genomic DNA from *C. difficile* strain 630Δerm. The SPA tag was amplified from plasmid p210, which has the DNA sequence of the SPA tag and *phoZ* with its RBS was amplified from pMC358. The appropriate PCR products were then cloned into the PmeI site of the vector pMSR via NEBuilder HiFi DNA Assembly (New England Biolabs). Toxin-mediated allelic exchange was then carried out as described previously ^32^.

To build the transcriptional *phoZ* gene reporter fusions, the riboswitch Cdi1_3 and its promoter region, the riboswitch Cdi1_6 and its promoter region, the promoter region only of *CD2309* and the promoter and the 5’ untranslated region of *CD2309*, including the Cdi1_9 riboswitch and the downstream DNA region up to the RBS, were amplified by PCR and cloned using NEBuilder HiFi DNA Assembly (New England Biolabs) into the vector pMC358 ^68^ linearized by inverse PCR. To build the translational *phoZ* gene reporter fusion, the *CD2309* promoter, the Cdi1_9 riboswitch, the native RBS, and the start codon of the *CD2309* gene were cloned into the vector pMC358 linearized by inverse PCR with primers designed to remove the RBS and the start codon of *phoZ*.

For inducible expression of *CD1980.2*, the CDS, along with the Cdi1_6 riboswitch, was amplified by PCR and cloned into SacI and BamHI sites of the pDIA6103 plasmid. The resulting pDIA6106 plasmid was then introduced into the required *C. difficile* strains. An inverse PCR approach was used to construct *CD1980.2*-*HA*-expressing plasmid (pDIA6574) on the basis of pDIA6106 with primers designed to introduce the hemagglutinin HA-tag sequence at the 3’ extremity of the CDS, directly upstream of the stop codon.

### Whole genome sequencing

The Δ7 mutant and the 630Δ*erm* parental strains were cultured in TY medium for 24 hours. Cells were collected, and genomic DNA was extracted using the NucleoSpin Microbial DNA Mini kit (Macherey-Nagel). Whole-genome sequencing was performed at Plasmidsaurus (https://www.plasmidsaurus.com/) using Oxford Nanopore Technologies (ONT) long-read sequencing. Quality control of the sequences was conducted using FastQC and NanoPlot via Galaxy Europe, and adapters were trimmed with Porechop. The resulting reads were aligned to the reference genome using minimap2. Small insertions and deletions (less than 50 bp) and SNPs were detected using Clair3, followed by normalization with bcftools norm and filtering with SnpSift Filter. Larger indels were identified with CuteSV.

### RNA sequencing

Strains 630Δ*erm* P*_tet_*-*dccA* (chromosomal insertion of P*_tet_* upstream of *dccA*) and ΔCdi1_6-*CD1980.2*/pCdi1_6-*CD1980.2* and the corresponding control strains (630Δ*erm* and ΔCdi1_6-*CD1980.2*/p) were grown to the late exponential phase in TY in the presence of 250 ng/mL ATc. Transcriptome analysis for each condition was performed with four independent total RNA preparations carried out using methods described before ^69^. The RNA samples were first treated using an Epicenter Bacterial Ribo-Zero kit. This depleted ribosomal RNA fraction was used to construct cDNA libraries using a TruSeq Stranded Total RNA sample prep kit (Illumina). Libraries were then sequenced by Illumina NextSeq 500. Cleaned sequenced reads were aligned to the genome of *C. difficile* strain 630 for the mapping of the sequences using Bowtie2. DESeq2 was used to perform normalization and differential analysis using values of the respective control strains serving as a reference for reporting the expression data of the P*_tet_*-*dccA* and ΔCdi1_6-*CD1980.2*/pP_tet_-Cdi1_6-*CD1980.2* strains Genes were considered differentially expressed if they showed a log_2_ fold change of greater than 2 or less than −2 and an adjusted *P*-value (*q*-value) of ≤ 0.05.

In addition, transcript abundance in wild-type 630Δ*erm* grown for 16 h on 70:30 sporulation agar was assessed by RNA sequencing performed by Novogene. This dataset was generated from three independent biological replicates. Raw reads were mapped to the genome of *C. difficile* strain 630, and gene expression was evaluated based on read counts rather than differential expression analysis.

### qRT-PCR analysis

*C. difficile* was grown in TY or 70:30 medium as described above, or on 70:30 sporulation agar. Cells were harvested from liquid cultures after 4 h of growth and from sporulation agar after 16 h. RNA extraction was performed as described previously ^69^. cDNA synthesis and quantitative real-time PCR were carried out as previously described ^14^.

Gene expression levels were normalized to the housekeeping gene *dnaF*. Relative expression was calculated using the ΔΔC_t_ method, with the indicated control condition used as reference. Data are presented as fold change relative to control conditions.

### Cell lysis, fractionation and western-blotting

Whole cell lysates were prepared using a single freeze-thaw cycle ^67^. Cell pellets were harvested, frozen at −20 °C, then thawed and resuspended in PBS with 0.12 μg/mL DNase I to OD_600_ = 40, and incubated at 37 °C for 40 min. Culture supernatants were filtered (0.22-μm) and precipitated on ice with 10% TCA for 30 min. then washed twice with 90% cold acetone and resuspended in PBS to OD_600_ = 40. For fractionation, pellets were resuspended in phosphate-sucrose buffer (0.05 M Na₂HPO₄, pH 7.0, 0.5 M sucrose) to OD_600_ = 40 and treated with 30 μg/mL CD27L endolysin at 37°C for 1 h ^24^. Supernatants (cell wall fraction) were collected, and protoplasts were lysed in phosphate buffer (0.05 M HNa_2_PO_4_, pH 7.0) containing 0.12 μg/mL DNase I at OD_600_ = 40 at 37 °C for 45 min. Cytoplasmic fractions were collected, and membranes were resuspended in phosphate buffer with 1% SDS at OD_600_ = 40.

Protein samples were mixed with equal volumes of 2x Laemmli buffer containing 12% β-mercaptoethanol, separated by electrophoresis and stained with InstantBlue (Sigma-Aldrich) for loading and fractionation controls. Western blotting was performed with anti-HA antibodies (Osenses) or anti-FLAG M2 (Sigma-Aldrich) antibodies using standard methods.

### Measurement of intracellular c-di-GMP concentration

*C. difficile* strains were grown in the presence of 25 or 250 ng/mL of the inducer ATc until they reached an OD_600_ of ∼0.6. A 10-mL culture aliquot was harvested by centrifugation and washed once. Nucleotides were extracted from the pellet with methanol/acetonitrile/Milli-Q water (40:40:20), and the c-di-GMP was detected and quantified by LC-MS/MS as described previously ^70^. To normalize the LC-MS/MS data, a 1-mL aliquot of the same bacterial culture was collected and pelleted by centrifugation. Bacterial pellets were subjected to whole-cell lysis. The lysates were centrifuged, and the protein content of the supernatant was determined using a bicinchoninic acid (BCA) assay kit (Thermo Fisher Scientific). The c-di-GMP concentrations are presented as picomol c-di-GMP per milligram of *C. difficile* total proteins.

### Alkaline phosphatase assay

*C. difficile* strains containing the *phoZ* reporter fusions were grown in TY medium, supplemented with thiamphenicol (7.5 mg/L) and ATc (25 to 250 ng/mL), until they reached an OD_600_ of ∼0.5 and a 2-mL culture aliquot was harvested by centrifugation. Samples were stored at −20 °C and alkaline phosphatase assays were performed as detailed elsewhere ^68^, without the use of chloroform for cell lysis.

### Sporulation assays

For assays on solid medium, sporulation frequencies of *C. difficile* strains were determined using a protocol published elsewhere ^71^. Briefly, overnight cultures of *C. difficile* in BHIS with 0.1% taurocholate and 0.2% fructose (plus 7.5 µg/mL thiamphenicol when required) were diluted and grown to OD_600_ ∼0.5. A 250-µL aliquot was then spread onto 70:30 agar plates containing thiamphenicol and 100 ng/mL ATc when needed. After 24 h, cells were scraped and resuspended in PBS to OD_600_ = 1.

For assays in liquid 70:30 medium, overnight cultures in TY were used to inoculate liquid 70:30 medium at an initial OD_600_ of 0.05 with 250 ng/mL ATc and 7.5 µg/mL thiamphenicol. Cultures were then incubated for 24 h.

For both conditions, sporulation efficiency was quantified using an ethanol-resistance assay ^71^. Samples were divided for vegetative and ethanol-resistant counts. Vegetative cells were determined by serial dilution and plating on BHIS agar. Ethanol-resistant spores were enumerated after 15 min treatment with 57% ethanol, dilution in PBS with 0.1% taurocholate, and plating on BHIS agar with taurocholate. Sporulation frequency was calculated as the number of ethanol-resistant spores / (number of vegetative cells + number of ethanol-resistant spores).

### Conventional mouse model infection studies

*C. difficile* spore inoculums were generated by plating 200 μl of an overnight culture of *C. difficile* grown onto 70:30 agar plates and incubating them at 37°C for 7 days in an anaerobic chamber. Spores were then harvested in 2 mL of ice-cold sterile water and purified by centrifugation using a HistoDenz (Sigma-Aldrich) gradient ^72^.

Six-week-old conventional female C57BL/6J mice (Charles River, France) were housed in groups of 5 per cage and maintained under a 12-h light-dark cycle at Plateforme des Animaleries-Inserm U1016, Institut Cochin. After acclimation for a week, all animals were treated for 3 days with a mixture of antibiotics dissolved in drinking water consisting of kanamycin (0.4 mg/mL), gentamicin (0.035 mg/mL), colistin (850 U/mL), metronidazole (0.215 mg/mL), and vancomycin (0.045 mg/mL) (Sigma Aldrich, France). After 1 day of wash-out, mice received a single intraperitoneal injection of clindamycin (10 mg/kg) (Sigma Aldrich). Twenty-four hours later, groups of ten mice were challenged with 10^5^ wild-type or Δ7 mutant *C. difficile* spores by oral gavage. Clinical monitoring, including weight, animal behavior, and stool consistency, was performed as previously described^73^: daily until day 4, and then every 3 days until day 18. To assess bacterial persistence, fecal pellets were collected over a 18-day period. Fecal pellets were homogenized in 1× prereduced PBS at a concentration of 50 mg/mL, serially diluted, and plated on chromID *C. difficile* agar (Biomerieux) to determine total CFU counts. Animals were euthanized on day 18 by intraperitoneal anesthesia (xylazine 10 mg/kg, ketamine 150 mg/kg), followed by cervical dislocation.

## ACKNOWLEDGEMENTS

This work was supported by grants from the French National Research Agency (ANR-20-CE15-0003-DIFFICROSS to J.P., ANR-13-JSV3-0005-01- CloSTARn, and ANR-22-CE15-0020-01- CdiffRib to O.S.), the Institut Universitaire de France (to O.S.), the University Paris-Saclay, the Institute for Integrative Biology of the Cell and the DIM-1HEALTH regional Ile-de-France programme (RPH21003DDP). We thank Johanne Delannoy for her technical assistance with the animal experiments and the members of the RNACLO lab for helpful discussions.

## AUTHOR CONTRIBUTIONS

Conceptualization: OS, JP. Data curation: JP. Funding acquisition: FB, JP. Investigation: ABL, LP, CR. Methodology: AJWD, RF, FB, OS, JP. Visualization: ABL, LP, JP. Funding acquisition: JP, BD, OS. Project administration: JP, BD. Supervision: JP, BD. Writing original draft: ABL, JP. Writing review & editing: ABL, AJWD, LP, RS, CR, FB, OS, JP.

## Data and materials availability

RNA-seq datasets generated in this study have been deposited in the NCBI Gene Expression Omnibus (GEO) under accession number GSE310232, including raw and processed data files. Additional RNA-seq data from wild-type 630Δ*erm* grown on 70:30 sporulation agar (16 h), as well as whole-genome sequencing data, have been deposited in the NCBI Sequence Read Archive (SRA) under BioProject accession PRJNA1363095. All other data needed to evaluate the conclusions are present in the paper or the Extended Data.

## Ethics statement

Animal studies were performed in agreement with European and French guidelines (Directive 86/609/CEE and Decree 87-848 of 19 October 1987). All animal procedures were approved by the “Ministère de l’Enseignement Supérieur et de la Recherche” under APAFIS#7600-2019070815198826 v2.

