## Supplementary information for "A family of small proteins links c-di-GMP riboswitch signaling with sporulation in *Clostridioides difficile*"

Supplementary Figure 1. RNA-seq visualization of the *dccA–CD1419* locus.

Supplementary Figure 2. A remnant ORF associated with the c-di-GMP-responding riboswitch Cdi1_7 encodes a truncated peptide homologous to CD1980.2.

Supplementary Figure 3. RgaR regulates three CD1980.2 orthologs not linked to a c-di-GMP riboswitch.

Supplementary Figure 4. Structural characterization of the riboswitch-associated small protein family.

Supplementary Figure 5. C-di-GMP requires the riboswitches to modulate the expression of the downstream genes.

Supplementary Figure 6. Sporulation-promoting activity of CD1980.2 in UK1 and stable expression of c-di-GMP riboswitch-associated genes under sporulation conditions.

Supplementary Figure 7. PCR verification of the seven deletions in the in *C. difficile* 630Δ*erm* Δ7 mutant.

Supplementary Figure 8. Low-abundance small protein paralogs contribute to sporulation in *Clostridioides difficile*.

Supplementary Table 1: Genes differentially expressed in the $P_{tet}$-*dccA* strain in comparison to the wild-type strain.

Supplementary Table 2. Sporulation-related genes downregulated in the $P_{tet}$-*dccA* strain in comparison to the wild-type strain.

Supplementary Table 3: Genes differentially expressed in the strain ΔCdi1_6–*CD1980.2* carrying the $P_{tet}$- Cdi1_6–*CD1980.2* plasmid in comparison to the same strain carrying an empty plasmid.

Supplementary Table 4: Sporulation-related genes upregulated in the strain ΔCdi1_6–*CD1980.2* carrying the $P_{tet}$- Cdi1_6–*CD1980.2* plasmid in comparison to the same strain carrying an empty plasmid.

Supplementary Table 5. Strains and plasmids used in this study.

Supplementary Table 6. Oligonucleotides used in this study.

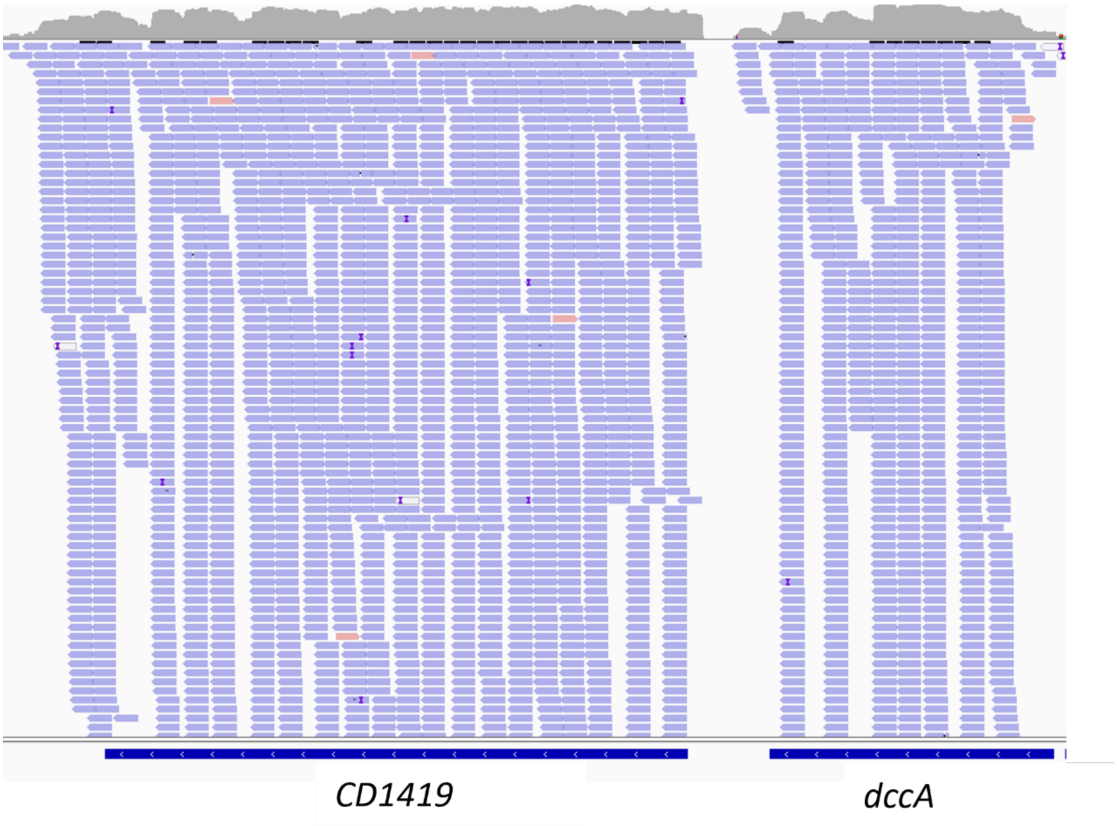


**Supplementary Figure 1:** RNA-seq visualization of the *dccA–CD1419* locus. RNA-seq reads from the $P_{tet}$-*dccA* strain were visualized using IGV. A representative genomic region spanning *dccA–CD1419* is shown. Reads aligned to the "+" strand are displayed in red and reads aligned to the "–" strand in blue.

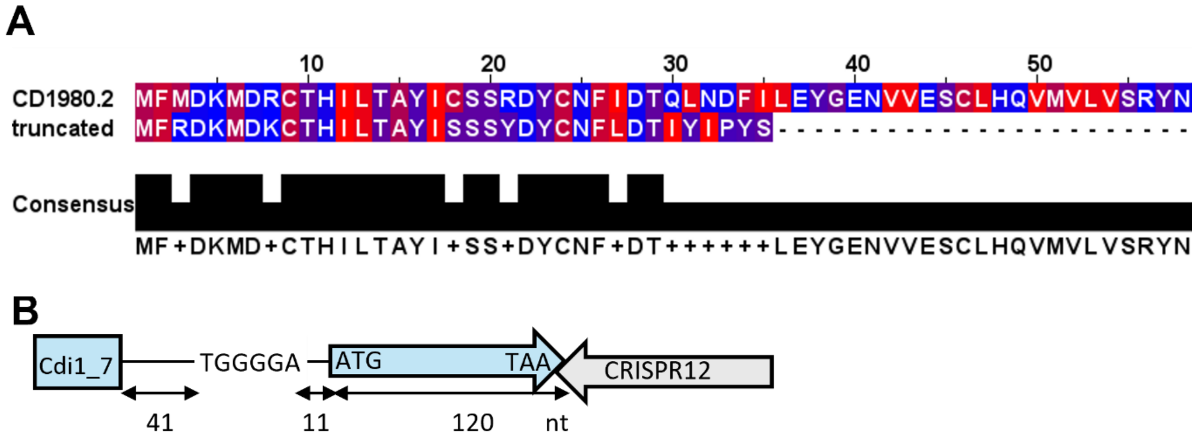


**Supplementary Figure 2. A remnant ORF associated with the c-di-GMP-responding riboswitch Cdi1_7 encodes a truncated peptide homologous to CD1980.2. (A)** Alignment of the small protein CD1980.2 with a truncated protein encoded by a remnant ORF associated with Cdi1_7. The hydrophobic and positively charged amino acids are indicated in red and blue, respectively. **(B)** Schematic representation of the Cdi1_7 riboswitch locus. The downstream remnant CDS is depicted as an arrow with the start (ATG) and the stop (TAA) codons, and the size of the CDS is indicated. The CRISPR array 12 located directly downstream disrupts the CDS. The putative RBS sequence (TGGGGA) is represented and the number of nucleotides between the RBS and the riboswitch and between the RBS and the CDS are indicated.

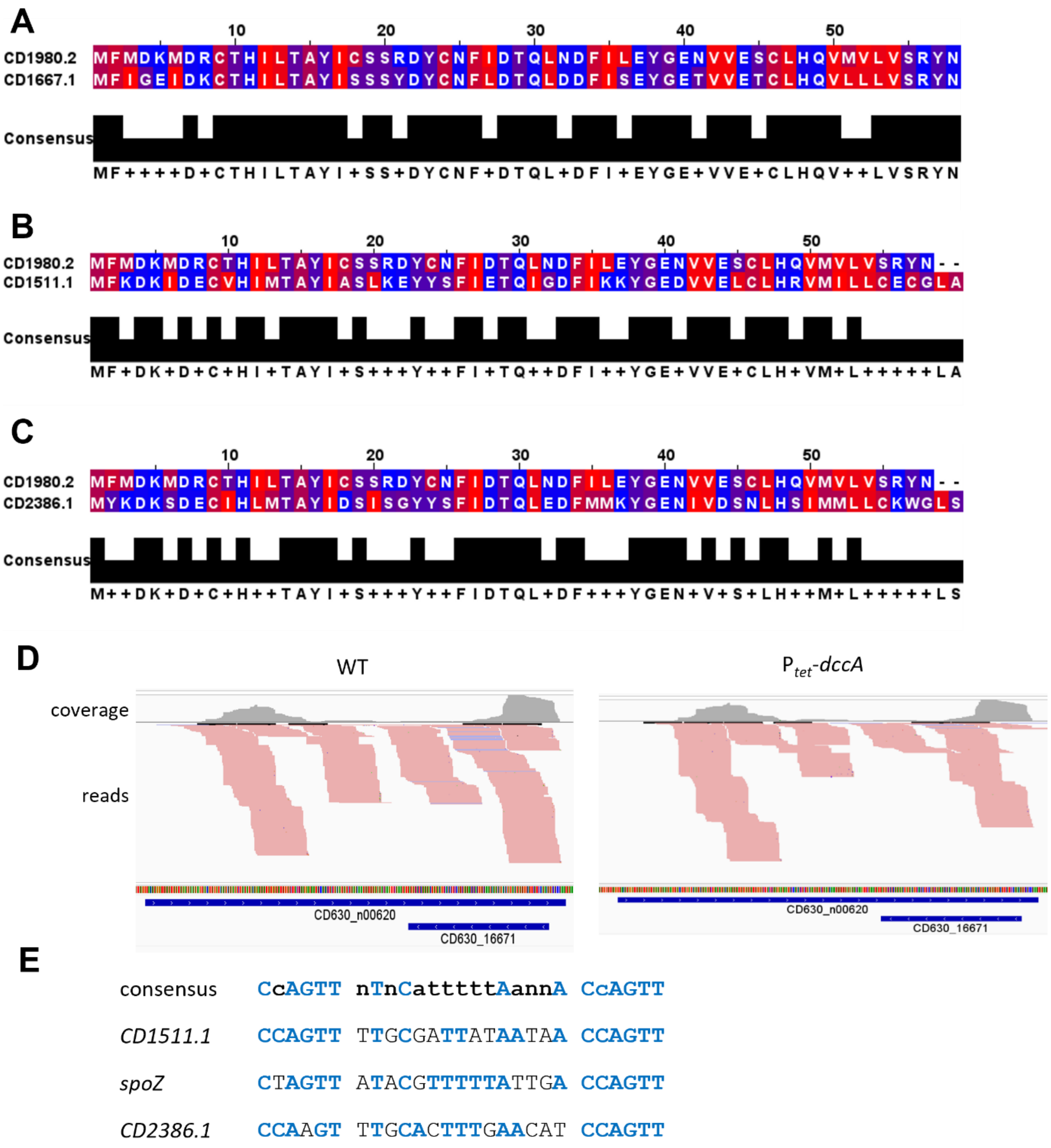


**Supplementary Figure 3. RgaR regulates three CD1980.2 orthologs not linked to a c-di-GMP riboswitch. (A to C)** Alignment of the small protein CD1980.2 with the small protein CD1667.1 **(A)**, CD1511.1 **(B)** or CD2386.1 **(C)**. The hydrophobic and positively charged amino acids are indicated in red and blue, respectively. **(D)** Visualization of RNA-seq data of the *CD1667.1*/*spoZ* (CD630_n00620) region from C. *difficile* wild type (WT) and $P_{tet}$-*dccA* , in which the diguanylate cyclase-encoding gene *dccA* is expressed from a chromosomally integrated anhydrotetracycline (ATc)-inducible promoter. Cultures for RNA-seq were grown in TY with 250 ng/ml ATc. "+" strand reads are shown in red, and "-" strand reads are shown in blue. Data are representative of four experiments. **(E)** Alignment of the putative RgaR binding sites with the promoters of *CD1511.1*, *spoZ* and *CD2386.1*.

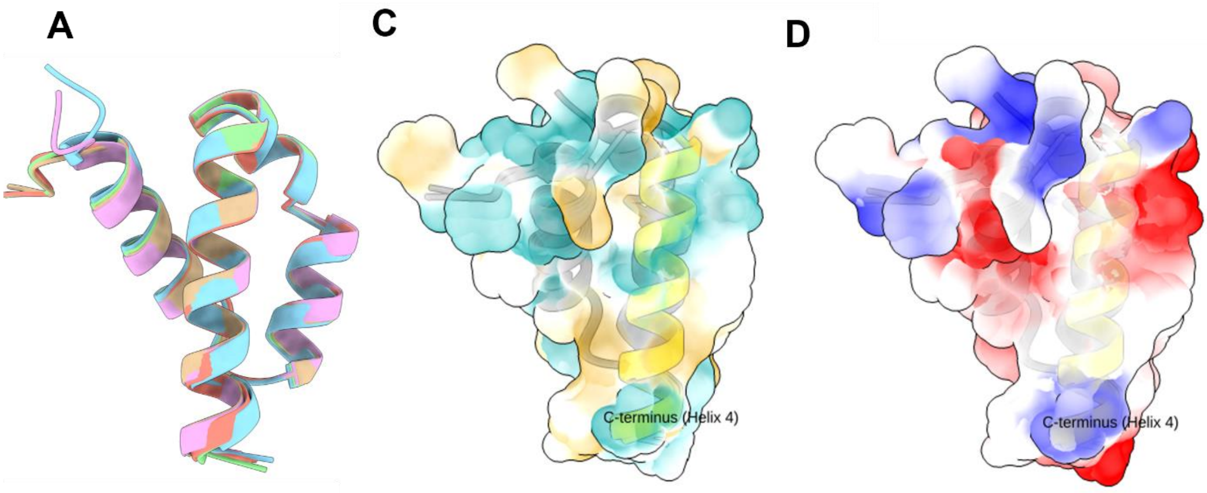


**B**

| Protein Model | H1 / H2 Angle (Oblique) | H1 / H3 Angle (Antiparallel) | H1 / H4 Angle (Anchor) | Anchor RMSD (Å) |
|---|---|---|---|---|
| Model 1 (CD2309) | 107.0° | 155.2° | 49.9° | 0.000 (Ref) |
| Model 2 (CD1424) | 113.3° | 143.4° | 37.4° | 0.192 |
| Model 3 (CD1980.2) | 96.8° | 159.3° | 61.6° | 0.223 |
| Model 4 (CD1990.3) | 94.0° | 152.9° | 63.9° | 0.216 |
| Model 5 (CD3368.2) | 106.1° | 155.1° | 49.9° | 0.215 |

**Supplementary Figure 4. Structural characterization of the riboswitch-associated small protein family. (A)** Structural superposition of AlphaFold3-predicted models of the five paralogs (CD2309, CD1424, CD1980_2, CD1990_3, and CD3368_2 demonstrating a conserved global architecture (global RMSD < 0.5 Å). Protein backbones are shown as cartoon representations, illustrating the strong structural similarity across paralogs despite moderate sequence divergence. (**B**) Quantitative interhelical angles and C-terminal anchor RMSD values for the five superposed models, derived from vector-based helical axis analysis. Helices 1 and 3 adopt an antiparallel arrangement (143.4–159.3°); Helices 2 and 4 adopt oblique orientations. Anchor RMSD values (0.19–0.22 Å) confirm exceptional geometric conservation of Helix 4 across all paralogs. (**C**) Molecular lipophilicity potential (MLP) mapped onto the solvent-excluded surfaces of the five superposed models. Yellow, hydrophobic; cyan, hydrophilic. The C-terminal Helix 4 (residues 40–57), rendered as a yellow cartoon, consistently exhibits a pronounced outward-facing hydrophobic surface patch across all models, supporting its proposed role as a membrane anchor. (**D**) Electrostatic surface potential of the four-helix bundle reveals a putative interaction interface. The solvent-excluded surface of the representative model (CD2309) is colored according to Coulombic electrostatic potential (blue, positive; red, negative; white, neutral). A prominent electropositive patch is located on the exposed surface of the helical core (Helices 1–3), suggesting a potential binding interface for negatively charged ligands or protein partners. In contrast, the C-terminal helix (Helix 4) is largely electrostatically neutral, consistent with its lipophilic character and role in membrane association.  Molecular graphics were generated using UCSF ChimeraX.

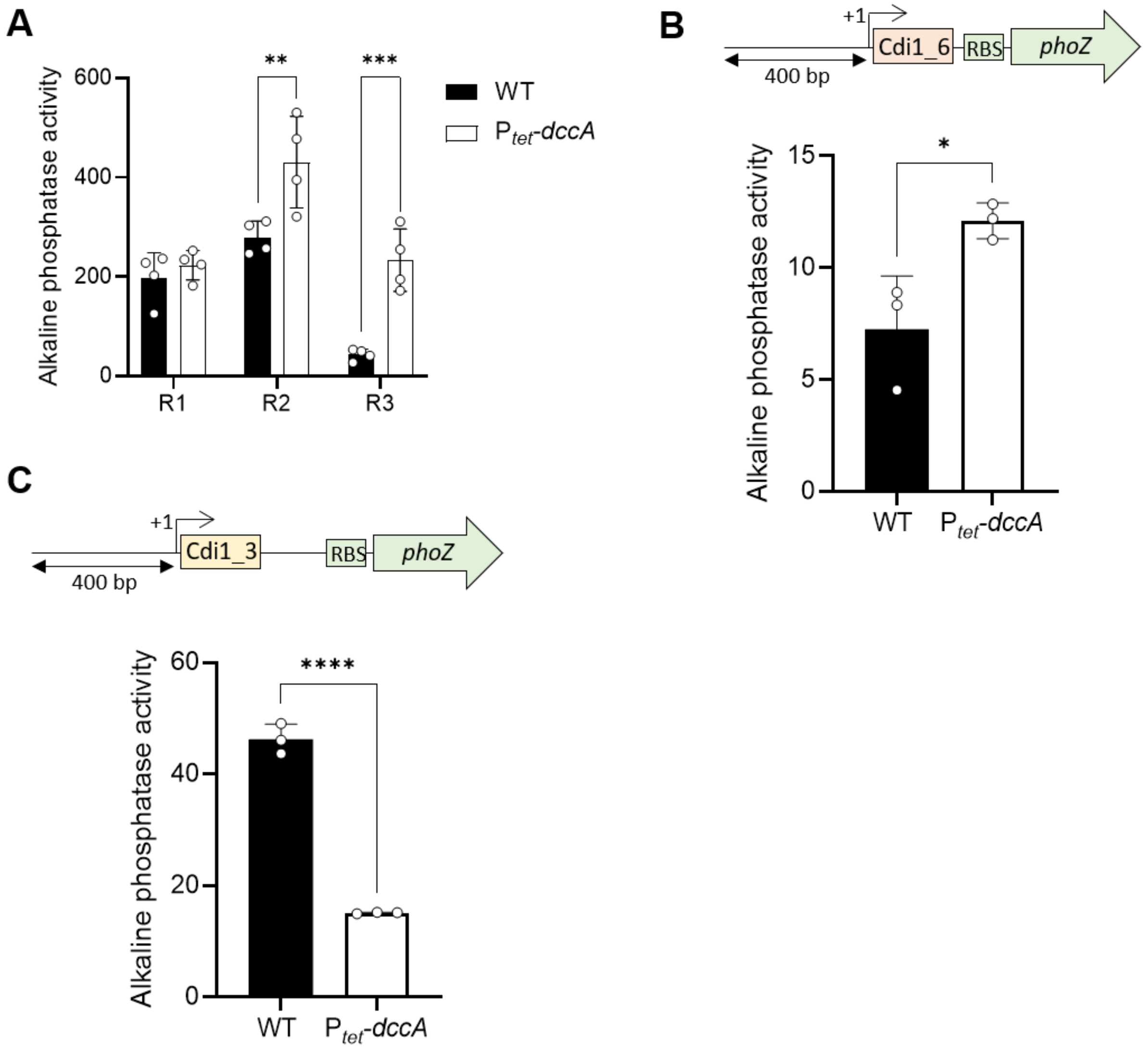


**Supplementary Figure 5. C-di-GMP requires the riboswitches to modulate the expression of the downstream genes. (A)** Alkaline phosphatase activity of the Cdi1_9::*phoZ* reporter fusions of various lengths (as depicted in Fig. 4A) in the wild-type (WT) and the $P_{tet}$-*dccA* strains, in which the diguanylate cyclase-encoding gene *dccA* is expressed from a chromosomally integrated anhydrotetracycline (ATc)-inducible promoter. Strains were grown in the exponential phase in TY with 25 ng/ml ATc. Bars represent means ± SD ($n = 4$ independent experiments). **P ≤ 0.01 and ***P ≤ 0.001 by a two-way ANOVA followed by a Sidak's multiple comparisons test. **(B)** Alkaline phosphatase activity of the Cdi1_6::*phoZ* reporter fusion in the WT and the $P_{tet}$-*dccA* strains. Strains were grown in the exponential phase in TY with 250 ng/ml ATc. A schematic of the Cdi1_6::*phoZ* transcriptional fusion carried on a plasmid is shown. Bars represent means ± SD ($n = 3$ independent experiments). *P ≤ 0.05 by an unpaired Student's *t* test. **(C)** Alkaline phosphatase activity of the Cdi1_3::*phoZ* reporter fusion in the WT and the $P_{tet}$-*dccA* strains. Strains were grown in the exponential phase in TY with 250 ng/ml ATc. A schematic of the Cdi1_3::*phoZ* transcriptional fusion carried on a plasmid is shown. Bars represent means ± SD ($n = 3$ independent experiments). ****P ≤ 0.0001 by an unpaired Student's *t* test.

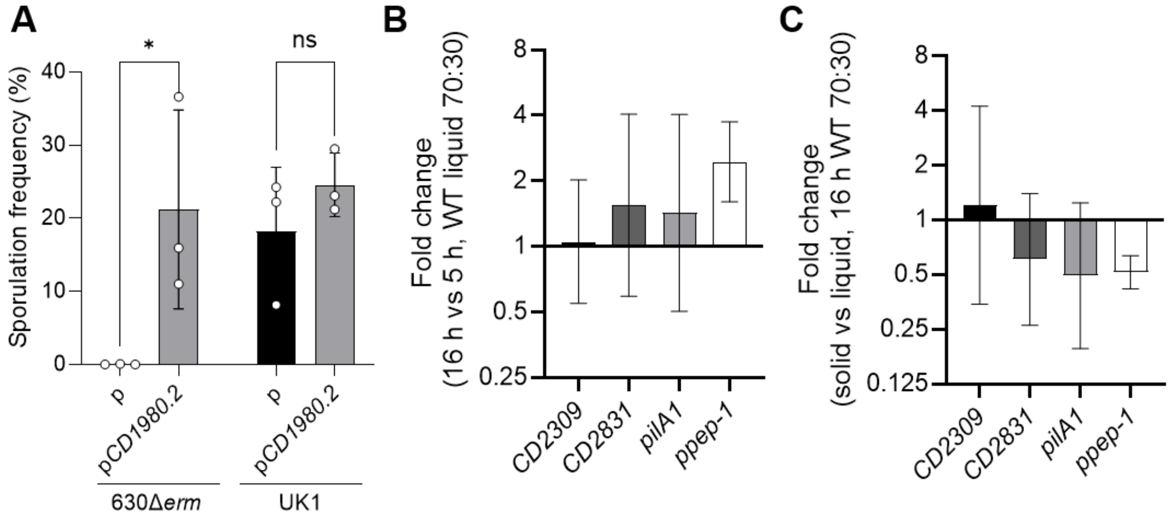


**Supplementary Figure 6. Sporulation-promoting activity of CD1980.2 in UK1 and stable expression of c-di-GMP riboswitch-associated genes under sporulation conditions. (A)** Overexpression of *CD1980.2* promotes sporulation in 630Δ*erm* but not in UK1 in liquid 70:30 medium. Ethanol-resistant spore formation of *C. difficile* 630Δ*erm*, and UK1 carrying an empty plasmid (p) or a plasmid expressing Cdi1_6-*CD1980.2* under the control of the $P_{tet}$ promoter (p*CD1980.2*) grown for 24h in liquid 70:30 medium in the presence of 250 ng/mL ATc. Bars represent means ± SD ($n$ = 3 independent experiments). *$P \leq 0.05$ by a two-way ANOVA followed by a Sidak's multiple comparison test. **(B)** and **(C)** Expression analysis of representative c-di-GMP riboswitch-associated genes during growth under sporulation conditions. Transcript levels of *CD2309*, *CD2831*, *pilA1* and *ppep-1* were measured by qRT-PCR in the wild-type strain grown in 70:30 medium. **(B)** Comparison of transcript levels between 4 h and 16 h in liquid culture. **(C)** Comparison of transcript levels between liquid and solid (70:30 agar) cultures after 16 h of growth. Bars represent means ± SD (n = 3 independent experiments).

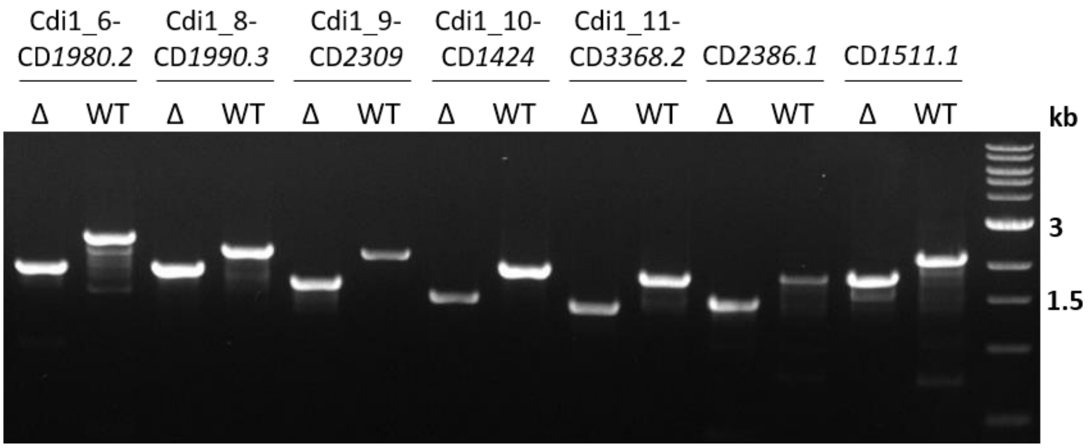


**Supplementary Figure 7. PCR verification of the seven deletions in the *C. difficile* 630Δ*erm* Δ7 mutant.** PCR amplification from wild type (WT) and Δ7 (Δ) to verify the indicated alleles.

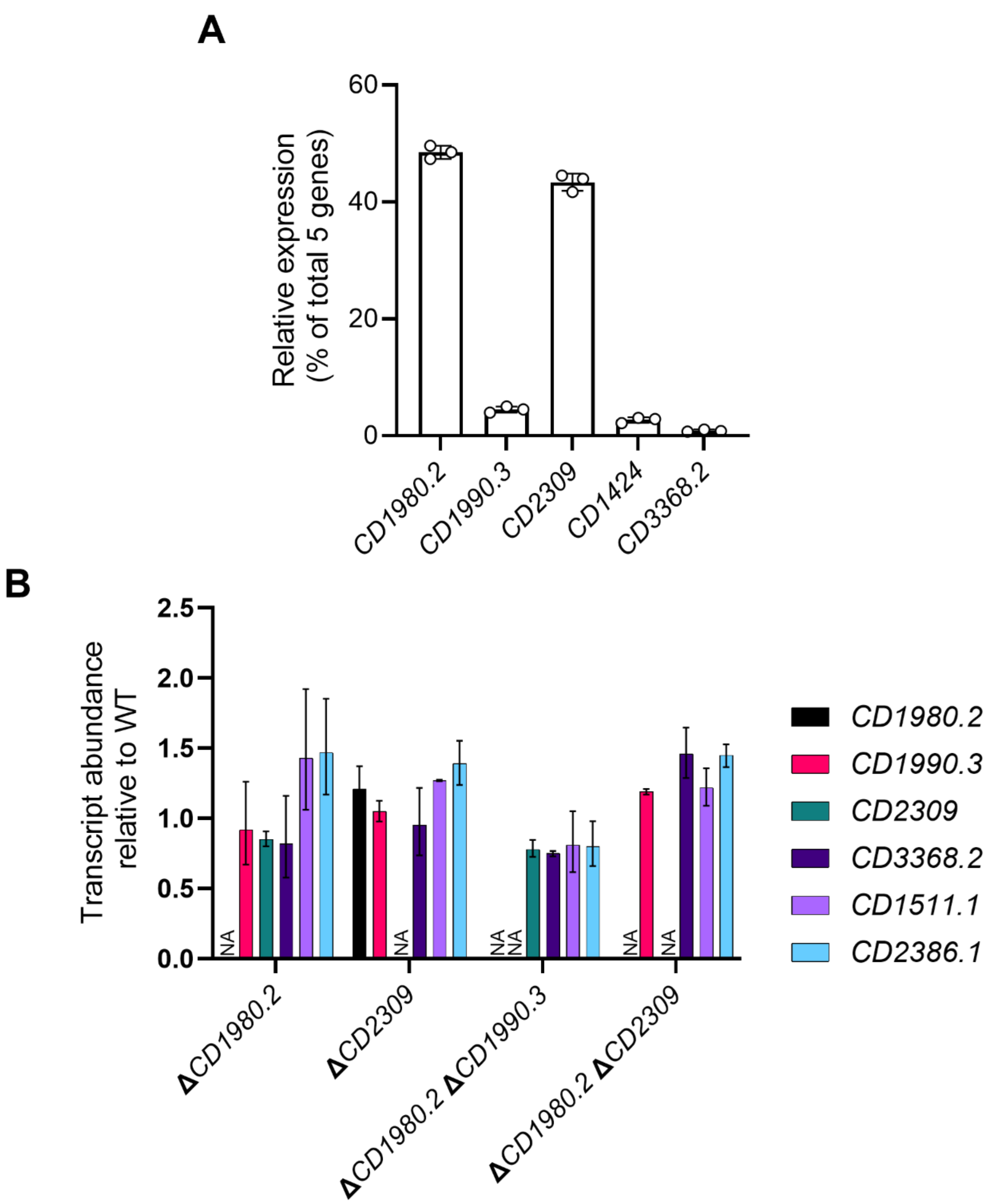


**Supplementary Figure 8. Low-abundance small protein paralogs contribute to sporulation in *Clostridioides difficile*.** **(A)** Relative mRNA abundance of the five small protein-encoding genes estimated from RNA-seq data obtained from the WT strain grown for 16 h on 70:30 sporulation agar supplemented with 250 ng/mL ATc. Bars represent means ± SD (n = 3 independent experiments). **(B)** qRT-PCR analysis showing expression levels of selected paralogs in 630Δ*erm* wild type and indicated deletion mutants grown in the exponential phase in TY medium. Bars represent means ± SD (n = 2 independent experiments). NA, not applicable, indicating absence of amplification for the deleted target gene in the corresponding mutant strain (Ct > 30).

**Supplementary Table 1: Genes differentially expressed in the $P_{tet}$-*dccA* strain in comparison to the wild-type strain.**

| Locus tag | Gene | $Log_2$ FC[a] | *q* value[b] | Annotated function |
|---|---|---|---|---|
| CD630_14200 | *dccA* | 7.745 | 8.69E-87 | diguanylate kinase signaling protein |
| CD630_14190 | | 4.643 | 1.64E-58 | putative diguanylate kinase signaling protein |
| CD630_19900 | | 3.748 | 4.58E-12 | putative protein with SH3 domain |
| CD630_35130 | | 3.726 | 1.18E-66 | putative pilin protein |
| CD630_28310 | | 3.463 | 6.60E-63 | putative adhesin |
| CD630_17160 | | 3.325 | 5.56E-05 | putative permease |
| CD630_Cdi1_1 | | 3.323 | 6.29E-03 | ncRNA |
| CD630_00801 | *rpmC* | 3.131 | 8.64E-09 | 50S ribosomal protein L29 |
| CD630_26730 | *appB* | 3.119 | 3.17E-07 | ABC-type transport system, oligopeptide-family permease protein |
| CD630_15800 | *hom2* | 3.007 | 3.87E-09 | Homoserine dehydrogenase |
| CD630_30160 | | 2.979 | 1.51E-04 | Transcription antiterminator, PTS operon regulator |
| CD630_25480 | | 2.883 | 1.74E-04 | ABC-type transport system, sugar-family permease |
| CD630_26720 | *appA* | 2.829 | 2.90E-07 | ABC-type transport system, oligopeptide-family solute-binding protein |
| CD630_25470 | | 2.791 | 3.36E-05 | conserved hypothetical protein |
| CD630_26710 | | 2.724 | 1.35E-03 | ABC-type transport system, ATP-binding protein; putative oligopeptide transport system |
| CD630_30150 | | 2.715 | 1.80E-04 | PTS system, mannose-specfic IIA component |
| CD630_25490 | | 2.714 | 1.05E-03 | ABC-type transport system, sugar-family permease |
| CD630_00820 | *rplN* | 2.705 | 6.95E-05 | 50S ribosomal protein L14 |
| CD630_00810 | *rpsQ* | 2.701 | 3.58E-05 | 30S ribosomal protein S17 |
| CD630_16840 | | 2.698 | 1.46E-08 | putative radical SAM superfamily protein |
| CD630_17170 | | 2.692 | 2.29E-04 | uncharacterised protein |
| CD630_26740 | *appC* | 2.645 | 3.36E-05 | ABC-type transport system, oligopeptide-family permease protein |
| CD630_00840 | *rplE* | 2.609 | 1.95E-05 | 50S ribosomal protein L5 |
| CD630_RNA_8 | | 2.562 | 3.85E-02 | ncRNA |
| CD630_n00290 | | 2.524 | 2.50E-03 | ncRNA |
| CD630_17151 | | 2.506 | 6.62E-03 | conserved hypothetical protein |
| CD630_00770 | *rpsS* | 2.488 | 1.01E-04 | 30S ribosomal protein S19 |
| CD630_00640 | *rplL* | 2.475 | 2.10E-06 | 50S ribosomal protein L7/L12 |
| CD630_30120 | | 2.412 | 5.91E-03 | putative alpha-mannosidase |
| CD630_n00830 | | 2.4 | 1.52E-04 | ncRNA |
| CD630_17680 | | 2.345 | 4.03E-04 | putative membrane protein |
| CD630_06730 | | 2.321 | 4.05E-05 | putative methyltransferase |
| CD630_21740 | | 2.288 | 6.24E-04 | ABC-type transport system,cystine/aminoacid-family extracellular solute-binding protein |
| CD630_21720 | | 2.287 | 1.26E-06 | ABC-type transport system,cystine/aminoacid-family ATP-binding protein |
| CD630_23270 | | 2.273 | 1.36E-02 | PTS system, fructose/mannitol family IIA component |
| CD630_00830 | *rplX* | 2.271 | 1.05E-03 | 50S ribosomal protein L24 |
| CD630_21750 | | 2.247 | 6.90E-04 | ABC-type transport system,cystine/aminoacid-family permease |
| CD630_32310 | *hpt1* | 2.22 | 3.94E-06 | Hypoxanthine phosphoribosyltransferase |
| CD630_07860 | | 2.208 | 1.34E-03 | putative membrane protein |
| CD630_31580 | | 2.206 | 5.03E-05 | Transcriptional regulator, TRAP family |
| CD630_21730 | | 2.198 | 4.89E-04 | putative peptidase, M20D family |
| CD630_08730 | | 2.158 | 2.38E-03 | ABC-type transport system, sugar-family extracellular solute-binding protein |
| CD630_21760 | | 2.146 | 1.11E-03 | ABC-type transport system,cystine/aminoacid-family permease |

| | | | | |
|---|---|---|---|---|
| CD630_21770 | | 2.116 | 2.05E-03 | ABC-type transport system,cystine/aminoacid-family extracellular solute-binding protein |
| CD630_30130 | | 2.113 | 4.43E-03 | PTS system, mannose-specfic IIC component |
| CD630_16160 | | 2.09 | 4.63E-06 | putative diguanylate kinase signaling protein |
| CD630_23050 | | 2.082 | 1.38E-05 | putative pilin protein |
| CD630_00800 | *rplP* | 2.074 | 1.06E-02 | 50S ribosomal protein L16 |
| CD630_07590 | *plfB* | 2.055 | 8.04E-04 | Formate acetyltransferase (Pyruvate formate-lyase) |
| CD630_23420 | *sucD* | 2.048 | 1.37E-20 | Succinate-semialdehyde dehydrogenase (NAD(P)+) |
| CD630_08740 | | 2.047 | 1.01E-04 | ABC-type transport system, sugar-family ATP-binding protein |
| CD630_25050 | | 2.02 | 6.04E-03 | Transcriptional regulator, TetR family |
| CD630_23550 | *trxA2* | -2.008 | 5.23E-04 | Thioredoxin 2 (Trx2) |
| CD630_15110 | *cotB* | -2.015 | 1.20E-02 | spore coat protein |
| CD630_32480 | | -2.026 | 9.40E-03 | Polysaccharide deacetylase |
| CD630_10340 | | -2.033 | 1.30E-03 | putative mannosyl-glycoprotein endo-beta-N-acetylglucosamidase |
| CD630_04460 | *oraE* | -2.041 | 3.56E-04 | D-ornithine aminomutase E component |
| CD630_28330 | | -2.042 | 9.03E-03 | putative calcium-transporting ATPase |
| CD630_34990 | *spoVT* | -2.042 | 9.35E-03 | Stage V sporulation protein T |
| CD630_10660 | | -2.039 | 2.11E-02 | conserved hypothetical protein |
| CD630_20800 | | -2.046 | 9.88E-03 | putative FAD-binding subunit of xanthine dehydrogenase |
| CD630_00220 | *fusA1* | -2.064 | 1.56E-05 | Elongation factor G (EF-G) |
| CD630_23470 | *proP* | -2.071 | 3.07E-10 | putative Xaa-Pro dipeptidase |
| CD630_11510 | *minE* | -2.071 | 6.29E-05 | Cell division topological specificity factor |
| CD630_20000 | *isp* | -2.074 | 1.42E-02 | Intracellular serine protease |
| CD630_13190 | | -2.076 | 4.58E-03 | putative polysaccharide deacetylase |
| CD630_16600 | | -2.083 | 4.37E-09 | uncharacterised protein |
| CD630_17410 | | -2.084 | 1.38E-04 | Fragment of selenoprotein B,glycine/betaine/sarcosine/D-proline reductase family (sarcosine reductase) |
| CD630_26510 | *murG* | -2.088 | 1.62E-04 | UDP-N-acetylglucosamine--N-acetylmuramyl-(pentapeptide) pyrophosphoryl-undecaprenol N-acetylglucosamine transferase |
| CD630_35220 | | -2.105 | 3.64E-02 | uncharacterised protein |
| CD630_15111 | | -2.113 | 1.84E-03 | conserved hypothetical protein |
| CD630_20980 | | -2.118 | 1.91E-05 | conserved hypothetical protein |
| CD630_10320 | | -2.133 | 2.09E-03 | conserved hypothetical protein |
| CD630_12910 | *dacF* | -2.133 | 7.99E-03 | D-alanyl-D-alanine carboxypeptidase (penicillin-binding protein) |
| CD630_24591 | | -2.138 | 2.16E-08 | conserved hypothetical protein |
| CD630_26250 | | -2.138 | 3.03E-07 | putative membrane protein |
| CD630_10330 | *mnaA* | -2.14 | 5.55E-04 | UDP-N-acetylglucosamine 2-epimerase (UDP-GlcNAc-2-epimerase) |
| CD630_18580 | | -2.141 | 2.91E-03 | putative cell surface protein Tn1549-like,CTn5-Orf15 |
| CD630_17400 | | -2.144 | 2.55E-04 | Glycine/sarcosine/betaine reductase complex,protein B, alpha and beta subunits |
| CD630_02360 | *fliS2* | -2.158 | 5.59E-05 | Flagellar protein FliS2 |
| CD630_03140 | | -2.16 | 4.85E-02 | putative membrane protein |
| CD630_19290 | | -2.178 | 1.04E-02 | putative membrane protein |
| CD630_19921 | | -2.186 | 5.16E-04 | conserved hypothetical protein |
| CD630_35640 | *spoIIR* | -2.185 | 9.57E-03 | Pro-sigma(E) endopeptidase (stage II sporulation) |
| CD630_36520 | | -2.182 | 1.93E-02 | putative peptidase, M1 family |
| CD630_26520 | *spoVE* | -2.198 | 1.02E-04 | Cell division/stage V sporulation protein |
| CD630_13540 | | -2.197 | 3.63E-02 | putative exported protein |
| CD630_23540 | *grdE* | -2.215 | 6.83E-05 | Glycine reductase complex component B subunits alpha and beta (Selenoprotein PB alpha/beta) [Contains: Betaine reductase |

| | | | | |
|---|---|---|---|---|
| | | | | component B subunit beta; Betaine reductase component B subunit alpha] |
| CD630_10631 | | -2.218 | 3.74E-02 | uncharacterised protein |
| CD630_06220 | | -2.238 | 4.61E-02 | conserved hypothetical protein |
| CD630_02310 | *flgK* | -2.245 | 1.03E-05 | Flagellar hook-associated protein FlgK (or HAP1) |
| CD630_28000 | | -2.243 | 2.44E-04 | putative membrane protein |
| CD630_30430 | | -2.251 | 3.59E-24 | conserved hypothetical protein |
| CD630_02370 | *fliD* | -2.269 | 7.45E-06 | Flagellar hook-associated protein 2 FliD (or HAP2) |
| CD630_01520 | *gcp* | -2.283 | 1.29E-06 | putative O-sialoglycoprotein endopeptidase |
| CD630_17890 | | -2.313 | 2.17E-02 | putative membrane protein |
| CD630_11490 | *minC* | -2.339 | 8.36E-07 | Cell division regulator (septum placement) |
| CD630_01310 | | -2.336 | 1.32E-03 | putative membrane protein |
| CD630_22451 | | -2.333 | 1.32E-03 | uncharacterised protein |
| CD630_26350 | | -2.337 | 1.94E-03 | putative spore envelope protein |
| CD630_14860 | | -2.335 | 4.51E-03 | putative lipoprotein |
| CD630_32960 | | -2.344 | 1.02E-05 | putative pilus assembly ATPase |
| CD630_12140 | *spo0A* | -2.35 | 1.23E-07 | Stage 0 sporulation protein A |
| CD630_04440 | *ortB* | -2.353 | 6.29E-05 | 2-amino-4-ketopentanoate thiolase beta subunit |
| CD630_23870 | | -2.355 | 2.13E-21 | Alpha-hydroxy acid dehydrogenase,FMN-dependent |
| CD630_31450 | | -2.356 | 6.46E-07 | putative serine-aspartate repeat-containing protein SdrF |
| CD630_11500 | *minD* | -2.384 | 1.04E-06 | Septum site-determining protein MinD (Cell division inhibitor MinD) |
| CD630_04400 | *cwp27* | -2.398 | 1.39E-11 | putative cell wall binding protein cwp27 |
| CD630_11730 | | -2.393 | 1.66E-10 | putative FAD-linked oxidase |
| CD630_11700 | | -2.396 | 2.65E-09 | conserved hypothetical protein |
| CD630_29660 | *adhE1* | -2.398 | 2.27E-06 | Aldehyde-alcohol dehydrogenase |
| CD630_02350 | *fliS1* | -2.396 | 2.24E-05 | Flagellar protein FliS1 |
| CD630_12590 | *brnQ-1* | -2.398 | 1.92E-04 | Branched chain amino acid transport system carrier protein |
| CD630_01500 | | -2.414 | 1.06E-05 | putative peptidase, M22 family |
| CD630_04470 | | -2.414 | 8.92E-04 | putative reactivating factor for adenosylcobalamine-dependent D-ornithine aminomutase |
| CD630_32890 | | -2.409 | 4.31E-03 | conserved hypothetical protein |
| CD630_14160 | | -2.413 | 2.65E-02 | putative membrane protein |
| CD630_30230 | | -2.415 | 1.32E-03 | conserved hypothetical protein |
| CD630_15431 | | -2.424 | 2.53E-08 | conserved hypothetical protein |
| CD630_04420 | *ord* | -2.434 | 4.05E-05 | 2,4-diaminopentanoate dehydrogenase |
| CD630_17260 | | -2.431 | 3.39E-03 | conserved hypothetical protein |
| CD630_11720 | *etfA4* | -2.445 | 5.07E-12 | Electron transfer flavoprotein subunit alpha |
| CD630_01250 | | -2.441 | 8.57E-05 | putative cell wall endopeptidase |
| CD630_06630 | *tcdA* | -2.451 | 2.44E-10 | Toxin A |
| CD630_03720 | | -2.447 | 1.07E-03 | putative cell wall hydrolase Tn916-like,CTn1-Orf17 |
| CD630_05710 | | -2.477 | 1.10E-04 | uncharacterised protein |
| CD630_02290 | *flgM* | -2.48 | 5.59E-05 | Negative regulator of flagellin synthesis (Anti-sigma-d factor) |
| CD630_26480 | | -2.502 | 3.43E-07 | conserved hypothetical protein |
| CD630_02320 | *flgL* | -2.508 | 7.10E-06 | Flagellar hook-associated protein FlgL (or HAP3) |
| CD630_26290 | *spoIVA* | -2.515 | 3.95E-04 | Stage IV sporulation protein A |
| CD630_32950 | | -2.542 | 2.09E-03 | putative type II secretion system protein |
| CD630_01260 | *spoIIID* | -2.537 | 2.97E-03 | Stage III sporulation protein D |
| CD630_35420 | *spmA* | -2.549 | 2.39E-02 | Spore maturation protein A |
| CD630_04430 | *ortA* | -2.575 | 1.16E-02 | 2-amino-4-ketopentanoate thiolase alpha subunit |

| | | | | |
|---|---|---|---|---|
| CD630_01510 | *rimI* | -2.603 | 5.86E-07 | putative alanine acetyltransferase, ribosomal family |
| CD630_32970 | | -2.597 | 1.53E-04 | conserved hypothetical protein |
| CD630_23520 | *grdA* | -2.612 | 2.46E-05 | Glycine reductase complex selenoprotein A (selenocysteine) |
| CD630_23750 | | -2.609 | 9.88E-03 | uncharacterised protein |
| CD630_07740 | *spoVAD* | -2.619 | 1.15E-03 | Stage V sporulation protein AD |
| CD630_33120 | | -2.617 | 3.23E-02 | Transporter, Major Facilitator Superfamily (MFS) |
| CD630_21270 | | -2.638 | 2.07E-17 | putative exported protein |
| CD630_05720 | | -2.638 | 6.95E-04 | putative sporulation protein |
| CD630_17880 | | -2.646 | 3.68E-02 | putative membrane protein |
| CD630_11710 | *etfB4* | -2.696 | 5.63E-11 | Electron transfer flavoprotein subunit alpha |
| CD630_03730 | | -2.697 | 8.66E-10 | putative membrane protein Tn916-like,CTn1-Orf18 |
| CD630_23740 | | -2.701 | 8.57E-03 | conserved hypothetical protein |
| CD630_11980 | *spoIIIAG* | -2.705 | 6.25E-04 | Stage III sporulation protein AG |
| CD630_10632 | | -2.707 | 1.66E-02 | conserved hypothetical protein |
| CD630_22620 | | -2.721 | 6.78E-10 | Transporter, Major Facilitator Superfamily (MFS) |
| CD630_26360 | | -2.722 | 7.52E-03 | putative membrane protein |
| CD630_24690 | *spoIIP* | -2.759 | 5.17E-04 | Stage II sporulation protein P |
| CD630_19000 | | -2.776 | 7.31E-06 | conserved hypothetical protein |
| CD630_10630 | | -2.794 | 4.05E-03 | conserved hypothetical protein |
| CD630_32920 | | -2.818 | 2.58E-03 | conserved hypothetical protein |
| CD630_02340 | *csrA* | -2.855 | 2.00E-05 | Carbon storage regulator homolog CsrA |
| CD630_04480 | *orr* | -2.854 | 6.45E-03 | ornithine racemase |
| CD630_11560 | | -2.909 | 7.09E-26 | conserved hypothetical protein |
| CD630_11960 | *spoIIIAE* | -2.907 | 3.84E-03 | Stage III sporulation protein AE |
| CD630_27620 | *uppS1* | -2.922 | 5.22E-08 | putative undecaprenyl pyrophosphate synthetase |
| CD630_11990 | *spoIIIAH* | -2.917 | 2.14E-04 | Stage III sporulation protein AH |
| CD630_24450 | | -2.931 | 1.48E-02 | putative transmembrane signaling protein,TspO/MBR family |
| CD630_05510 | *sleC* | -2.942 | 1.98E-02 | Spore cortex-lytic enzyme pre-pro-form |
| CD630_04450 | *oraS* | -2.977 | 2.77E-02 | D-ornithine aminomutase S component |
| CD630_21120 | | -3.003 | 6.54E-06 | uncharacterised protein |
| CD630_23510 | *grdB* | -3.017 | 1.49E-07 | Glycine reductase complex component B gamma subunit (selenocysteine) |
| CD630_26490 | | -3.039 | 1.11E-07 | conserved hypothetical protein |
| CD630_35630 | | -3.069 | 1.87E-07 | putative spore cortex-lytic hydrolase |
| CD630_11970 | *spoiIIIAF* | -3.075 | 4.27E-04 | Stage III sporulation protein AF |
| CD630_02390 | *fliC* | -3.097 | 1.04E-07 | Flagellin C |
| CD630_11940 | *spoIIIAC* | -3.114 | 6.29E-04 | Stage III sporulation protein AC |
| CD630_01490 | | -3.117 | 7.42E-07 | putative atp/gtp hydrolase |
| CD630_22150 | | -3.126 | 4.27E-04 | Transcriptional regulator, HTH-type |
| CD630_33660 | | -3.137 | 5.37E-36 | putative exported protein |
| CD630_24700 | *gpr* | -3.129 | 3.01E-05 | Spore endopeptidase |
| CD630_SQ1376 | | -3.14 | 4.13E-05 | ncRNA |
| CD630_27370 | | -3.16 | 1.80E-11 | putative nitrilase/cyanide hydratase and apolipoprotein N-acyltransferase |
| CD630_11920 | *spoIIIAA* | -3.159 | 2.48E-05 | Stage III sporulation protein AA |
| CD630_19350 | *spoVS* | -3.174 | 3.13E-09 | Stage V sporulation protein S |
| CD630_27380 | | -3.197 | 3.29E-13 | putative cytosine permease |
| CD630_32900 | | -3.204 | 5.93E-05 | conserved hypothetical protein |
| CD630_11950 | *spoIIIAD* | -3.216 | 7.16E-04 | Stage III sporulation protein AD |
| CD630_15670 | *cotG* | -3.243 | 3.15E-03 | spore coat protein-manganese catalase |

| | | | | |
|---|---|---|---|---|
| CD630_07790 | | -3.254 | 5.74E-11 | putative amidohydrolase, M20D peptidase family |
| CD630_14300 | | -3.247 | 2.44E-04 | putative delta-lactam-biosynthetic de-N-acteylase |
| CD630_33670 | | -3.252 | 8.18E-04 | conserved hypothetical protein |
| CD630_12720 | | -3.264 | 7.79E-06 | putative magnesium chelatase |
| CD630_24780 | | -3.284 | 4.71E-04 | putative DNA uptake transporter |
| CD630_24680 | | -3.283 | 8.07E-03 | conserved hypothetical protein |
| CD630_32940 | | -3.297 | 4.62E-06 | putative type IV pilin PilA |
| CD630_30240 | | -3.29 | 1.22E-04 | uncharacterised protein |
| CD630_07780 | | -3.305 | 3.04E-08 | conserved hypothetical protein |
| CD630_11930 | *spoIIIAB* | -3.316 | 2.82E-03 | Stage III sporulation protein AB |
| CD630_07800 | | -3.33 | 3.87E-11 | conserved hypothetical protein, DUF1177 family |
| CD630_07770 | | -3.349 | 4.21E-11 | putative membrane protein |
| CD630_06840 | | -3.369 | 1.84E-07 | putative ATP-dependent peptidase, M41 family |
| CD630_19802 | | -3.403 | 7.41E-14 | conserved hypothetical protein |
| CD630_12980 | | -3.432 | 1.83E-09 | putative sporulation protein |
| CD630_23861 | | -3.425 | 1.57E-04 | conserved hypothetical protein |
| CD630_35511 | | -3.441 | 3.77E-03 | putative membrane protein |
| CD630_n01110 | | -3.464 | 5.11E-03 | ncRNA |
| CD630_22140 | *SinR* | -3.487 | 1.06E-05 | Transcriptional regulator, HTH-type |
| CD630_23730 | | -3.529 | 1.09E-07 | putative CstA-like carbon starvation protein |
| CD630_12440 | | -3.567 | 5.38E-05 | conserved hypothetical protein |
| CD630_28080 | | -3.619 | 1.09E-07 | conserved hypothetical protein |
| CD630_32930 | | -3.634 | 2.82E-03 | putative pilus assembly protein |
| CD630_12450 | | -3.736 | 2.47E-04 | conserved hypothetical protein |
| CD630_07710 | *spoIIAB* | -3.779 | 3.07E-10 | Anti-sigma F factor (Stage II sporulation protein AB) |
| CD630_07720 | *sigF* | -3.875 | 6.42E-09 | RNA polymerase sigma-F factor |
| CD630_07700 | *spoIIAA* | -3.909 | 4.21E-08 | Anti-sigma F factor antagonist |
| CD630_19410 | | -3.97 | 1.10E-05 | uncharacterised protein |
| CD630_Cdi1_6 | | -4.002 | 3.78E-13 | ncRNA |
| CD630_35512 | | -4.019 | 1.87E-06 | conserved hypothetical protein |
| CD630_26871 | | -4.052 | 2.59E-02 | uncharacterised protein |
| CD630_26420 | *sigG* | -4.112 | 1.82E-08 | RNA polymerase sigma-G factor |
| CD630_26560 | *spoVD* | -4.186 | 4.21E-11 | Stage V sporulation protein D (Sporulation-specific penicillin-binding protein) |
| CD630_19670 | | -4.217 | 1.50E-07 | uncharacterised protein |
| CD630_Cdi1_10 | | -4.258 | 1.18E-21 | ncRNA |
| CD630_01480 | | -4.291 | 3.87E-11 | uncharacterised protein |
| CD630_25622 | | -4.323 | 2.77E-10 | uncharacterised protein |
| CD630_14240 | | -4.406 | 5.39E-20 | conserved hypothetical protein |
| CD630_12430 | | -4.409 | 7.78E-07 | conserved hypothetical protein |
| CD630_26500 | | -4.458 | 7.27E-07 | putative cell division protein Fts-Q type |
| CD630_26430 | *sigE* | -4.459 | 7.13E-09 | RNA polymerase sigma-E factor |
| CD630_26570 | | -4.504 | 1.56E-08 | conserved hypothetical protein |
| CD630_24970 | *comE* | -4.58 | 2.14E-08 | Competence protein ComEA |
| CD630_34890 | | -4.693 | 4.65E-08 | putative oligoendopeptidase F, peptidase M3B family |
| CD630_12710 | | -4.724 | 1.20E-03 | putative restriction endonuclease |
| CD630_Cdi1_11 | | -4.768 | 6.43E-15 | ncRNA |
| CD630_33682 | | -4.856 | 2.42E-19 | conserved hypothetical protein |
| CD630_26440 | *spoIIGA* | -4.857 | 1.19E-08 | Sporulation sigma-E factor processing peptidase |

| | | | | |
|---|---|---|---|---|
| CD630_28090 | | -4.891 | 3.94E-06 | uncharacterised protein |
| CD630_34900 | *spoIIE* | -4.938 | 4.21E-11 | stage II sporulation protein |
| CD630_12420 | | -5.28 | 8.10E-03 | conserved hypothetical protein |
| CD630_23090 | | -5.436 | 2.96E-36 | conserved hypothetical protein |
| CD630_SQ2503 | | -5.428 | 1.07E-03 | ncRNA |
| CD630_Cdi1_9 | | -5.66 | 5.37E-36 | ncRNA |
| CD630_Cdi1_8 | | -5.752 | 2.63E-25 | ncRNA |
| CD630_19903 | | -5.685 | 7.24E-18 | conserved hypothetical protein |
| CD630_25630 | | -6.466 | 4.55E-04 | conserved hypothetical protein |

[a]The $\log_2$ fold changes listed are averages from four biological replicates.
[b]The *q* value is an adjusted *P* value, taking into account the false discovery rate. The *q* values listed are the averages from four biological replicates.

**Supplementary Table 2. Sporulation-related genes downregulated in the $P_{tet}$-*dccA* strain in comparison to the wild-type strain.**

| Locus tag | Gene | $Log_2$ FC[b] | *q* value[c] | Annotated function |
|---|---|---|---|---|
| **dependent on Spo0A[a]** | | | | |
| CD630_05710 | | -2.477 | 0.00010992 | uncharacterised protein |
| CD630_05720 | | -2.638 | 0.00069497 | putative sporulation protein |
| CD630_06220 | | -2.238 | 0.04613819 | conserved hypothetical protein |
| CD630_07700 | *spoIIAA* | -3.909 | 4.2108E-08 | Anti-sigma F factor antagonist |
| CD630_07710 | *spoIIAB* | -3.779 | 3.0672E-10 | Anti-sigma F factor (Stage II sporulation protein AB) |
| CD630_07720 | *sigF* | -3.875 | 6.4211E-09 | RNA polymerase sigma-F factor |
| CD630_10330 | *mnaA* | -2.14 | 0.00055483 | UDP-N-acetylglucosamine 2-epimerase (UDP-GlcNAc-2-epimerase) |
| CD630_11700 | | -2.396 | 2.6494E-09 | conserved hypothetical protein |
| CD630_12140 | *spo0A* | -2.35 | 1.23E-07 | Stage 0 sporulation protein A |
| CD630_19410 | | -3.97 | 1.1028E-05 | uncharacterised protein |
| CD630_19670 | | -4.217 | 1.4956E-07 | uncharacterised protein |
| CD630_23730 | | -3.529 | 1.0908E-07 | putative CstA-like carbon starvation protein |
| CD630_23740 | | -2.701 | 0.00856768 | conserved hypothetical protein |
| CD630_26420 | *sigG* | -4.112 | 1.8229E-08 | RNA polymerase sigma-G factor |
| CD630_26430 | *sigE* | -4.459 | 7.1266E-09 | RNA polymerase sigma-E factor |
| CD630_26440 | *spoIIGA* | -4.857 | 1.1896E-08 | Sporulation sigma-E factor processing peptidase |
| CD630_26510 | *murG* | -2.088 | 0.00016196 | UDP-N-acetylglucosamine--N-acetylmuramyl-(pentapeptide) pyrophosphoryl-undecaprenol N-acetylglucosamine transferase |
| CD630_26520 | *spoVE* | -2.198 | 0.00010233 | Cell division/stage V sporulation protein |
| CD630_26560 | *spoVD* | -4.186 | 4.208E-11 | Stage V sporulation protein D (Sporulation-specific penicillin-binding protein) |
| CD630_26570 | | -4.504 | 1.5571E-08 | conserved hypothetical protein |
| CD630_32890 | | -2.409 | 0.00431074 | conserved hypothetical protein |
| CD630_32900 | | -3.204 | 5.9304E-05 | conserved hypothetical protein |
| CD630_34890 | | -4.693 | 4.6535E-08 | putative oligoendopeptidase F, peptidase M3B family |
| CD630_34900 | *spoIIE* | -4.938 | 4.208E-11 | stage II sporulation protein |
| CD630_35630 | | -3.069 | 1.8728E-07 | putative spore cortex-lytic hydrolase |
| CD630_35640 | *spoIIR* | -2.185 | 0.00956647 | Pro-sigma(E) endopeptidase (stage II sporulation) |
| **dependent on $\sigma^E$[a]** | | | | |
| CD630_01260 | *spoIIID* | -2.537 | 0.00296831 | Stage III sporulation protein D |
| CD630_01310 | | -2.336 | 0.00131542 | putative membrane protein |
| CD630_10630 | | -2.794 | 0.00405312 | conserved hypothetical protein |
| CD630_10660 | | -2.039 | 0.02112493 | conserved hypothetical protein |
| CD630_11920 | *spoIIIAA* | -3.159 | 2.4769E-05 | Stage III sporulation protein AA |
| CD630_11930 | *spoIIIAB* | -3.316 | 0.00281548 | Stage III sporulation protein AB |
| CD630_11940 | *spoIIIAC* | -3.114 | 0.00062856 | Stage III sporulation protein AC |
| CD630_11950 | *spoIIIAD* | -3.216 | 0.00071603 | Stage III sporulation protein AD |
| CD630_11960 | *spoIIIAE* | -2.907 | 0.0038387 | Stage III sporulation protein AE |
| CD630_11970 | *spoiIIIAF* | -3.075 | 0.00042738 | Stage III sporulation protein AF |
| CD630_11980 | *spoIIIAG* | -2.705 | 0.00062474 | Stage III sporulation protein AG |
| CD630_11990 | *spoIIIAH* | -2.917 | 0.00021379 | Stage III sporulation protein AH |
| CD630_12590 | *brnQ-1* | -2.398 | 0.00019215 | Branched chain amino acid transport system carrier protein |
| CD630_13190 | | -2.076 | 0.00457591 | putative polysaccharide deacetylase |
| CD630_15110 | *cotB* | -2.015 | 0.01204859 | spore coat protein |
| CD630_17260 | | -2.431 | 0.00338684 | conserved hypothetical protein |

| | | | | |
|---|---|---|---|---|
| CD630_17400 | | -2.144 | 0.00025513 | Glycine/sarcosine/betaine reductase complex,protein B, alpha and beta subunits |
| CD630_17410 | | -2.084 | 0.00013797 | Fragment of selenoprotein B,glycine/betaine/sarcosine/D-proline reductase family (sarcosine reductase) |
| CD630_19290 | | -2.178 | 0.01040488 | putative membrane protein |
| CD630_20000 | *isp* | -2.074 | 0.014215 | Intracellular serine protease |
| CD630_24450 | | -2.931 | 0.0147539 | putative transmembrane signaling protein,TspO/MBR family |
| CD630_26290 | *spoIVA* | -2.515 | 0.00039501 | Stage IV sporulation protein A |
| CD630_28000 | | -2.243 | 0.00024376 | putative membrane protein |
| CD630_28330 | | -2.042 | 0.00903349 | putative calcium-transporting ATPase |
| CD630_32480 | | -2.026 | 0.00940261 | Polysaccharide deacetylase |
| CD630_35220 | | -2.105 | 0.03644994 | uncharacterised protein |
| CD630_35420 | *spmA* | -2.549 | 0.02389208 | Spore maturation protein A |
| CD630_35512 | | -4.019 | 1.8677E-06 | conserved hypothetical protein |
| CD630_36520 | | -2.182 | 0.01932694 | putative peptidase, M1 family |
| **dependent on $\sigma^{F}$[a]** | | | | |
| CD630_01250 | | -2.441 | 8.5692E-05 | putative cell wall endopeptidase |
| CD630_22451 | | -2.333 | 0.00131542 | uncharacterised protein |
| CD630_23750 | | -2.609 | 0.00987615 | uncharacterised protein |
| CD630_24690 | *spoIIP* | -2.759 | 0.00051681 | Stage II sporulation protein P |
| CD630_24700 | *gpr* | -3.129 | 3.0147E-05 | Spore endopeptidase |
| CD630_27620 | *uppS1* | -2.922 | 5.2176E-08 | putative undecaprenyl pyrophosphate synthetase |
| **dependent on $\sigma^{G}$[a]** | | | | |
| CD630_06840 | | -3.369 | 1.8398E-07 | putative ATP-dependent peptidase, M41 family |
| CD630_07740 | *spoVAD* | -2.619 | 0.00115201 | Stage V sporulation protein AD |
| CD630_12910 | *dacF* | -2.133 | 0.00798899 | D-alanyl-D-alanine carboxypeptidase (penicillin-binding protein) |
| CD630_12980 | | -3.432 | 1.8278E-09 | putative sporulation protein |
| CD630_13540 | | -2.197 | 0.03626462 | putative exported protein |
| CD630_14300 | | -3.247 | 0.00024376 | putative delta-lactam-biosynthetic de-N-acteylase |
| CD630_14860 | | -2.335 | 0.00450586 | putative lipoprotein |
| CD630_15670 | *cotG* | -3.243 | 0.00314562 | spore coat protein-manganese catalase |
| CD630_21120 | | -3.003 | 6.5389E-06 | uncharacterised protein |
| CD630_26350 | | -2.337 | 0.00193732 | putative spore envelope protein |
| CD630_26360 | | -2.722 | 0.00751623 | putative membrane protein |
| CD630_26871 | | -4.052 | 0.0258766 | uncharacterised protein |
| CD630_28080 | | -3.619 | 1.0858E-07 | conserved hypothetical protein |
| CD630_28090 | | -4.891 | 3.9424E-06 | uncharacterised protein |
| CD630_33120 | | -2.617 | 0.03227951 | Transporter, Major Facilitator Superfamily (MFS) |
| CD630_34990 | *spoVT* | -2.042 | 0.00935224 | Stage V sporulation protein T |
| CD630_35511 | | -3.441 | 0.00376798 | putative membrane protein |
| **dependent on $\sigma^{K}$[a]** | | | | |
| CD630_05510 | *sleC* | -2.942 | 0.01981565 | Spore cortex-lytic enzyme pre-pro-form |
| CD630_10631 | | -2.218 | 0.03742119 | uncharacterised protein |
| CD630_10632 | | -2.707 | 0.01660459 | conserved hypothetical protein |

[a]The SpoA and sporulation-associated sigma factors regulons are based on those reported in [1].
[b]The $\log_2$ fold changes listed are averages from four biological replicates.
[c]The *q* value is an adjusted *P* value, taking into account the false discovery rate. The *q* values listed are the averages from four biological replicates.

**Supplementary Table 3: Genes differentially expressed in the strain ΔCdi1_6–*CD1980.2* carrying the $P_{tet}$- Cdi1_6–*CD1980.2* plasmid in comparison to the same strain carrying an empty plasmid.**

| Locus tag | Gene | $Log_2$ FC[a] | *q* value[b] | Annotated function |
|---|---|---|---|---|
| CD630_19802 | | 13.183 | 4.06E-32 | conserved hypothetical protein |
| CD630_Cdi1_6 | | 12.359 | 3.34E-29 | ncRNA |
| CD630_32490 | *sspB* | 8.029 | 7.96E-106 | Small, acid-soluble spore protein beta |
| CD630_24000 | *cotJB2* | 7.02 | 2.71E-29 | Spore coat peptide assembly protein CotJB 2 |
| CD630_02140 | | 6.86 | 1.66E-10 | uncharacterised protein |
| CD630_10632 | | 6.703 | 4.92E-82 | conserved hypothetical protein |
| CD630_24010 | *cotD* | 6.552 | 1.81E-24 | Spore coat protein, manganese catalase |
| CD630_23990 | | 6.372 | 6.69E-11 | putative spore coat protein |
| CD630_06190 | | 6.359 | 1.58E-46 | conserved hypothetical protein |
| CD630_10633 | | 6.211 | 5.18E-42 | conserved hypothetical protein |
| CD630_06220 | | 6.178 | 7.43E-73 | conserved hypothetical protein |
| CD630_34391 | | 6.082 | 1.41E-05 | uncharacterised protein |
| CD630_06200 | | 6.013 | 2.17E-32 | conserved hypothetical protein |
| CD630_07830 | *spoIVB2* | 5.709 | 1.88E-15 | Stage IV sporulation protein SpoIVB, S55 peptidase family |
| CD630_12300 | *sigK* | 5.604 | 3.85E-21 | Sigma K |
| CD630_02130 | | 5.553 | 3.53E-09 | putative spore coat protein |
| CD630_26880 | *sspA* | 5.366 | 1.47E-90 | Small, acid-soluble spore protein alpha |
| CD630_05510 | *sleC* | 5.3 | 1.03E-31 | Spore cortex-lytic enzyme pre-pro-form |
| CD630_06210 | | 5.157 | 8.41E-17 | putative membrane protein |
| CD630_SQ995 | *RCd23* | 5.149 | 1.31E-03 | ncRNA |
| CD630_01290 | | 5.04 | 2.84E-29 | putative sporulation protein yyac |
| CD630_33500 | | 4.998 | 1.55E-09 | putative glycosyl transferase, family 2 |
| CD630_10670 | *CdeC* | 4.937 | 2.14E-43 | exosporium cysteine rich protein |
| CD630_14330 | *cotE* | 4.882 | 7.96E-18 | spore coat protein: peroxiredoxin/chitinase |
| CD630_10631 | | 4.772 | 3.30E-34 | uncharacterised protein |
| CD630_05970 | *cotF* | 4.547 | 1.01E-06 | Spore coat peptide assembly protein CotF |
| CD630_05960 | | 4.424 | 9.88E-06 | uncharacterised protein |
| CD630_12910 | *dacF* | 4.377 | 2.86E-19 | D-alanyl-D-alanine carboxypeptidase (penicillin-binding protein) |
| CD630_35800 | | 4.371 | 4.82E-10 | uncharacterised protein |
| CD630_16310 | *sodA* | 4.35 | 1.05E-13 | spore coat protein-superoxide dismutase (Mn) |
| CD630_10650 | | 4.248 | 1.15E-26 | uncharacterised protein |
| CD630_21440 | | 4.226 | 1.37E-04 | putative membrane protein |
| CD630_03320 | *bclA1* | 4.188 | 4.78E-13 | putative exosporium glycoprotein |
| CD630_15810 | | 4.009 | 1.50E-06 | uncharacterised protein |
| CD630_33490 | *bclA3* | 3.999 | 7.39E-14 | Exosporium glycoprotein BclA3 |
| CD630_05980 | *cotCB* | 3.997 | 3.30E-05 | Spore-coat protein, manganese catalase |
| CD630_21120 | | 3.939 | 1.46E-18 | uncharacterised protein |
| CD630_20550 | | 3.929 | 4.46E-04 | conserved hypothetical protein |
| CD630_33120 | | 3.898 | 1.70E-07 | Transporter, Major Facilitator Superfamily (MFS) |
| CD630_30270 | | 3.843 | 1.37E-04 | PTS system, glucose-like IIA component |
| CD630_36130 | | 3.824 | 6.39E-05 | uncharacterised protein |
| CD630_30300 | | 3.807 | 2.02E-04 | PTS system, glucose-like IIBC component |
| CD630_19040 | | 3.795 | 7.78E-08 | ABC-type transport system, permease |
| CD630_16130 | *cotA* | 3.792 | 6.50E-07 | spore coat assembly protein |

| | | | | |
|---|---|---|---|---|
| CD630_30280 | | 3.775 | 1.74E-04 | putative phosphosugar isomerase |
| CD630_30320 | | 3.743 | 7.50E-06 | putative aminotransferase |
| CD630_14860 | | 3.685 | 4.05E-07 | putative lipoprotein |
| CD630_24090 | | 3.681 | 9.12E-14 | conserved hypothetical protein |
| CD630_18840 | | 3.662 | 8.91E-12 | conserved hypothetical protein |
| CD630_35511 | | 3.629 | 4.90E-12 | putative membrane protein |
| CD630_30290 | *malY* | 3.617 | 1.11E-03 | Bifunctional protein: cystathionine beta-lyase / repressor |
| CD630_32300 | *bclA2* | 3.588 | 3.94E-06 | putative exosporium glycoprotein |
| CD630_18450 | | 3.555 | 1.09E-11 | |
| CD630_22451 | | 3.453 | 1.09E-11 | uncharacterised protein |
| CD630_36200 | | 3.414 | 7.90E-05 | uncharacterised protein |
| CD630_17070 | | 3.358 | 4.74E-12 | putative C4-dicarboxylate anaerobic carrier,DcuC family |
| CD630_23430 | *cat1* | 3.351 | 1.08E-05 | Succinyl-CoA:coenzyme A transferase |
| CD630_18460 | | 3.234 | 7.46E-11 | putative conjugative transposon protein Tn1549-like, CTn5-Orf2 |
| CD630_30170 | | 3.216 | 5.91E-03 | putative glucose uptake protein |
| CD630_07930 | | 3.194 | 3.22E-03 | putative membrane protein, DUF81 family |
| CD630_07730 | *spoVAC* | 3.178 | 1.64E-06 | Stage V sporulation protein AC |
| CD630_30150 | | 3.122 | 4.77E-21 | PTS system, mannose-specfic IIA component |
| CD630_30140 | | 3.034 | 4.18E-37 | PTS system, mannose-specfic IIB component |
| CD630_13540 | | 3.017 | 3.62E-08 | putative exported protein |
| CD630_09020 | | 3.014 | 8.04E-05 | putative cation efflux protein |
| CD630_29680 | *spoVFA* | 2.965 | 3.37E-08 | Dipicolinate synthase subunit A |
| CD630_23260 | | 2.942 | 2.47E-02 | PTS system, fructose/mannitol family IIB component |
| CD630_23410 | *abfD* | 2.908 | 1.94E-05 | Gamma-aminobutyrate metabolism dehydratase/isomerase [includes: 4-hydroxybutyryl-coa dehydratase; vinylacetyl-coa-delta-isomerase] |
| CD630_30160 | | 2.903 | 8.94E-13 | Transcription antiterminator, PTS operon regulator |
| CD630_23400 | | 2.881 | 1.45E-05 | uncharacterised protein |
| CD630_30130 | | 2.836 | 2.05E-38 | PTS system, mannose-specfic IIC component |
| CD630_07740 | *spoVAD* | 2.825 | 4.41E-07 | Stage V sporulation protein AD |
| CD630_23440 | | 2.801 | 1.74E-04 | putative membrane protein |
| CD630_28450 | *rbr1* | 2.779 | 1.88E-09 | Rubrerythrin |
| CD630_23270 | | 2.764 | 2.90E-02 | PTS system, fructose/mannitol family IIA component |
| CD630_30120 | | 2.762 | 5.47E-12 | putative alpha-mannosidase |
| CD630_07750 | *spoVAE* | 2.747 | 5.70E-05 | Stage V sporulation protein AE |
| CD630_24310 | | 2.744 | 4.79E-04 | putative nitrite/sulphite reductase |
| CD630_21410 | | 2.717 | 2.78E-04 | Serine-type D-Ala-D-Ala carboxypeptidase |
| CD630_29341 | | 2.688 | 6.44E-03 | putative phage protein |
| CD630_25990 | | 2.684 | 1.14E-02 | putative transcriptional regulator |
| CD630_14300 | | 2.674 | 9.22E-10 | putative delta-lactam-biosynthetic de-N-acteylase |
| CD630_21210 | | 2.656 | 9.96E-07 | conserved hypothetical protein |
| CD630_29370 | | 2.642 | 1.92E-05 | putative phage protein |
| CD630_29350 | | 2.639 | 1.37E-02 | conserved hypothetical protein |
| CD630_23380 | *4hbD* | 2.63 | 4.85E-05 | 4-hydroxybutyrate dehydrogenase (4-hydroxybutanoate:NAD+ oxidoreductase) |
| CD630_06240 | | 2.596 | 2.15E-07 | putative transcriptional regulator, activator Mor |
| CD630_23390 | *cat2* | 2.553 | 1.28E-04 | 4-hydroxybutyrate CoA transferase |
| CD630_06750 | | 2.534 | 1.68E-03 | putative acetyltransferase |
| CD630_15670 | *cotG* | 2.485 | 1.45E-08 | spore coat protein-manganese catalase |
| CD630_23250 | | 2.471 | 1.38E-02 | PTS system, fructose/mannitol family IIC component |

| | | | | |
|---|---|---|---|---|
| CD630_29420 | | 2.442 | 1.12E-06 | putative phage resolvase/integrase |
| CD630_29670 | *spoVFB* | 2.422 | 4.48E-07 | Dipicolinate synthase subunit B |
| CD630_12301 | | 2.416 | 1.53E-08 | #N/A |
| CD630_23240 | | 2.369 | 1.54E-02 | putative sugar-phosphate dehydrogenase |
| CD630_25980 | | 2.363 | 4.61E-03 | putative oligosaccharide deacetylase |
| CD630_SQ2150 | | 2.356 | 1.00E-02 | ncRNA |
| CD630_16780 | | 2.336 | 2.03E-03 | putative membrane protein |
| CD630_31160 | *bglF3* | 2.335 | 1.66E-03 | PTS system, beta-glucoside-specific IIABC component, bglF3 |
| CD630_23750 | | 2.327 | 3.41E-05 | uncharacterised protein |
| CD630_07920 | | 2.294 | 8.99E-03 | putative membrane protein, DUF81 family |
| CD630_34990 | *spoVT* | 2.285 | 1.06E-07 | Stage V sporulation protein T |
| CD630_SQ476 | | 2.27 | 8.34E-04 | ncRNA |
| CD630_30360 | | 2.27 | 1.25E-02 | Transporter, Major Facilitator Superfamily (MFS) |
| CD630_30310 | | 2.247 | 1.45E-03 | Transcription antiterminator, PTS operon regulator |
| CD630_03110 | | 2.245 | 1.28E-07 | putative spore coat assembly asparagine-rich protein |
| CD630_29380 | | 2.24 | 4.43E-04 | putative phage protein |
| CD630_10660 | | 2.237 | 6.39E-08 | conserved hypothetical protein |
| CD630_13700 | | 2.209 | 9.09E-03 | putative phage XkdS-like protein |
| CD630_31360 | *bglA7* | 2.203 | 3.95E-02 | 6-phospho-beta-glucosidase, bglA7 |
| CD630_22140 | *SinR* | 2.129 | 2.14E-15 | Transcriptional regulator, HTH-type |
| CD630_29490 | | 2.127 | 6.41E-06 | Transcriptional regulator, Phage-type |
| CD630_23230 | | 2.126 | 2.22E-02 | putative sugar-phosphate dehydrogenase |
| CD630_25120 | | 2.101 | 2.47E-02 | PTS system, glucose-like IIA component |
| CD630_29480 | | 2.1 | 2.62E-07 | putative phage protein |
| CD630_26360 | | 2.099 | 4.90E-05 | putative membrane protein |
| CD630_23420 | *sucD* | 2.097 | 4.11E-04 | Succinate-semialdehyde dehydrogenase (NAD(P)+) |
| CD630_14630 | | 2.077 | 1.39E-13 | uncharacterised protein |
| CD630_12340 | | 2.072 | 4.30E-04 | putative phage protein |
| CD630_29440 | | 2.056 | 6.41E-09 | putative phage essential recombination function protein |
| CD630_29310 | | 2.025 | 3.14E-04 | putative phage endodeoxyribonuclease RusA-like |
| CD630_22160 | | -2.798 | 1.51E-07 | uncharacterised protein |
| CD630_n00330 | | -3.089 | 3.27E-02 | ncRNA |
| CD630_04900 | | -3.727 | 1.51E-07 | putative sugar-phosphate dehydrogenase |

[a]The $\log_2$ fold changes listed are averages from four biological replicates.
[b]The *q* value is an adjusted *P* value, taking into account the false discovery rate. The *q* values listed are the averages from four biological replicates.

**Supplementary Table 4: Sporulation-related genes upregulated in the strain ΔCdi1_6–*CD1980.2* carrying the $P_{tet}$- Cdi1_6–*CD1980.2* plasmid in comparison to the same strain carrying an empty plasmid.**

| Locus tag | Gene | $Log_2$ FC[b] | *q* value[c] | Annotated function |
|---|---|---|---|---|
| **dependent on Spo0A[a]** | | | | |
| CD630_06190 | | 6.359 | 1.5778E-46 | conserved hypothetical protein |
| CD630_06200 | | 6.013 | 2.1697E-32 | conserved hypothetical protein |
| CD630_06210 | | 5.157 | 8.4098E-17 | putative membrane protein |
| CD630_06220 | | 6.178 | 7.4334E-73 | conserved hypothetical protein |
| CD630_12340 | | 2.072 | 0.00043047 | putative phage protein |
| CD630_14630 | | 2.077 | 1.389E-13 | uncharacterised protein |
| CD630_15810 | | 4.009 | 1.4952E-06 | uncharacterised protein |
| CD630_16780 | | 2.336 | 0.00203089 | putative membrane protein |
| CD630_30320 | | 3.743 | 7.4959E-06 | putative aminotransferase |
| **dependent on $\sigma^{E}$[a]** | | | | |
| CD630_01290 | | 5.04 | 2.8368E-29 | putative sporulation protein yyac |
| CD630_02130 | | 5.553 | 3.5329E-09 | putative spore coat protein |
| CD630_03110 | | 2.245 | 1.2777E-07 | putative spore coat assembly asparagine-rich protein |
| CD630_05970 | *cotF* | 4.547 | 1.0079E-06 | Spore coat peptide assembly protein CotF |
| CD630_10660 | | 2.237 | 6.3907E-08 | conserved hypothetical protein |
| CD630_16130 | *cotA* | 3.792 | 6.4983E-07 | spore coat assembly protein |
| CD630_18450 | | 3.555 | 1.0915E-11 | putative membrane protein Tn1549-like,CTn5-Orf1 |
| CD630_18460 | | 3.234 | 7.4645E-11 | putative conjugative transposon protein Tn1549-like, CTn5-Orf2 |
| CD630_18840 | | 3.662 | 8.9128E-12 | conserved hypothetical protein |
| CD630_20550 | | 3.929 | 0.00044611 | conserved hypothetical protein |
| CD630_21210 | | 2.656 | 9.9618E-07 | conserved hypothetical protein |
| CD630_32300 | *bclA2* | 3.588 | 3.9365E-06 | putative exosporium glycoprotein |
| **dependent on $\sigma^{F}$[a]** | | | | |
| CD630_07830 | *spoIVB2* | 5.709 | 1.8764E-15 | Stage IV sporulation protein SpoIVB, S55 peptidase family |
| CD630_22451 | | 3.453 | 1.0915E-11 | uncharacterised protein |
| CD630_23750 | | 2.327 | 3.4136E-05 | uncharacterised protein |
| **dependent on $\sigma^{G}$[a]** | | | | |
| CD630_02140 | | 6.86 | 1.6642E-10 | uncharacterised protein |
| CD630_07730 | *spoVAC* | 3.178 | 1.6446E-06 | Stage V sporulation protein AC |
| CD630_07740 | *spoVAD* | 2.825 | 4.4073E-07 | Stage V sporulation protein AD |
| CD630_07930 | | 3.194 | 0.00322054 | putative membrane protein, DUF81 family |
| CD630_12910 | *dacF* | 4.377 | 2.8637E-19 | D-alanyl-D-alanine carboxypeptidase (penicillin-binding protein) |
| CD630_13540 | | 3.017 | 3.6152E-08 | putative exported protein |
| CD630_14300 | | 2.674 | 9.219E-10 | putative delta-lactam-biosynthetic de-N-acteylase |
| CD630_14860 | | 3.685 | 4.0542E-07 | putative lipoprotein |
| CD630_15670 | *cotG* | 2.485 | 1.4514E-08 | spore coat protein-manganese catalase |
| CD630_16310 | *sodA* | 4.35 | 1.049E-13 | spore coat protein-superoxide dismutase (Mn) |
| CD630_17070 | | 3.358 | 4.7351E-12 | putative C4-dicarboxylate anaerobic carrier,DcuC family |
| CD630_21120 | | 3.939 | 1.4647E-18 | uncharacterised protein |
| CD630_24310 | | 2.744 | 0.00047872 | putative nitrite/sulphite reductase |
| CD630_25980 | | 2.363 | 0.00460922 | putative oligosaccharide deacetylase |
| CD630_25990 | | 2.684 | 0.01138106 | putative transcriptional regulator |

| | | | | |
|---|---|---|---|---|
| CD630_26360 | | 2.099 | 4.8951E-05 | putative membrane protein |
| CD630_26880 | *sspA* | 5.366 | 1.4742E-90 | Small, acid-soluble spore protein alpha |
| CD630_28450 | *rbr1* | 2.779 | 1.8847E-09 | Rubrerythrin |
| CD630_32490 | *sspB* | 8.029 | 7.957E-106 | Small, acid-soluble spore protein beta |
| CD630_33120 | | 3.898 | 1.7013E-07 | Transporter, Major Facilitator Superfamily (MFS) |
| CD630_34990 | *spoVT* | 2.285 | 1.0561E-07 | Stage V sporulation protein T |
| CD630_35511 | | 3.629 | 4.8988E-12 | putative membrane protein |
| **dependent on $\sigma^{K}$[a]** | | | | |
| CD630_03320 | *bclA1* | 4.188 | 4.7817E-13 | putative exosporium glycoprotein |
| CD630_05510 | *sleC* | 5.3 | 1.032E-31 | Spore cortex-lytic enzyme pre-pro-form |
| CD630_05960 | | 4.424 | 9.8753E-06 | |
| CD630_05980 | *cotCB* | 3.997 | 3.295E-05 | Spore-coat protein, manganese catalase |
| CD630_09020 | | 3.014 | 8.0361E-05 | putative cation efflux protein |
| CD630_10631 | | 4.772 | 3.3015E-34 | uncharacterised protein |
| CD630_10632 | | 6.703 | 4.923E-82 | conserved hypothetical protein |
| CD630_10633 | | 6.211 | 5.1772E-42 | conserved hypothetical protein |
| CD630_10650 | | 4.248 | 1.1535E-26 | uncharacterised protein |
| CD630_10670 | *CdeC* | 4.937 | 2.1425E-43 | exosporium cysteine rich protein |
| CD630_12300 | *sigK* | 5.604 | 3.8495E-21 | Sigma K |
| CD630_14330 | *cotE* | 4.882 | 7.9585E-18 | spore coat protein: peroxiredoxin/chitinase |
| CD630_19040 | | 3.795 | 7.7844E-08 | ABC-type transport system, permease |
| CD630_21440 | | 4.226 | 0.00013664 | putative membrane protein |
| CD630_23990 | | 6.372 | 6.6924E-11 | putative spore coat protein |
| CD630_24000 | *cotJB2* | 7.02 | 2.71E-29 | Spore coat peptide assembly protein CotJB 2 |
| CD630_24010 | *cotD* | 6.552 | 1.8117E-24 | Spore coat protein, manganese catalase |
| CD630_24090 | | 3.681 | 9.1188E-14 | conserved hypothetical protein |
| CD630_29670 | *spoVFB* | 2.422 | 4.4844E-07 | Dipicolinate synthase subunit B |
| CD630_29680 | *spoVFA* | 2.965 | 3.373E-08 | Dipicolinate synthase subunit A |
| CD630_33490 | *bclA3* | 3.999 | 7.3921E-14 | Exosporium glycoprotein BclA3 |
| CD630_33500 | | 4.998 | 1.5534E-09 | putative glycosyl transferase, family 2 |
| CD630_35800 | | 4.371 | 4.8226E-10 | uncharacterised protein |

[a]The SpoA and sporulation-associated sigma factors regulons are based on those reported in [1].
[b]The $\log_2$ fold changes listed are averages from four biological replicates.
[c]The *q* value is an adjusted *P* value, taking into account the false discovery rate. The *q* values listed are the averages from four biological replicates.

**Supplementary Table 5. Strains and plasmids used in this study.**

| **Strain** | **Genotype** | **Origin** |
|---|---|---|
| ***E. coli*** | | |
| NEB-10 beta | Δ*(ara-leu) 7697 araD139 fhuA* Δ*lacX74 galK16 galE15 e14-* ϕ*80*d*lacZ*Δ*M15 recA1 relA1 endA1 nupG rpsL* ($Str^R$) *rph spoT1* Δ*(mrr-hsdRMS-mcrBC)* | New England Biolabs |
| HB101 (RP4) | *supE44 aa14 galK2 lacY1* Δ(*gpt-proA*) 62 *rpsL20 (*$Str^R$*)xyl-5 mtl-1 recA13* Δ(*mcrC-mrr*) *hsdS*$_B$ ($r_B$-$m_B$-) RP4 ($Tra^+$ IncP $Ap^R$ $Km^R$ $Tc^R$) | Laboratory stock |
| ***C. difficile*** | | |
| 630Δ*erm* | 630 Δ*ermB* | Laboratory stock [2] |
| CDIP369 (630/p) | 630 Δ*erm* strain carrying the empty plasmid pDIA6103 | Laboratory stock |
| CNRS_CD378 | 630Δ*erm* $P_{tet}$-*dccA*, chromosomally integrated ATc-inducible promoter driving *dccA* expression | This work |
| CDIP601 | 630Δ*erm* ΔCdi1_6-*CD1980.2* | This work |
| CDIP964 | CDIP601 carrying pDIA6574 for inducible Cdi1_6-*CD1980.2::HA* expression | This work |
| CNRS_CD497 | 630Δ*erm CD2309::SPA,* chromosomally integrated SPA tag fused in frame to the 3’ end of *CD2309* | This work |
| CNRS_CD727 | CNRS_CD497 carrying pDIA6103 | This work |
| CNRS_CD695 | CNRS_CD497 carrying pDIA5987 for inducible *dccA* expression | This work |
| CNRS_CD449 | 630Δ*erm* carrying p216 (region *CD2309::phoZ* reporter fusion) | This work |
| CNRS_CD467 | CNRS_CD378 carrying p216 (region 1 *CD2309::phoZ* reporter fusion | This work |
| CNRS_CD424 | 630Δ*erm* carrying p215 (region 2 *CD2309::phoZ* reporter fusion) | This work |
| CNRS_CD447 | CNRS_CD378 carrying p215 (region 2 *CD2309::phoZ* reporter fusion) | This work |

| | | |
|---|---|---|
| CNRS_CD468 | 630Δ*erm* carrying p230 (region 3 *CD2309::phoZ* reporter fusion) | This work |
| CNRS_CD470 | CNRS_CD378 carrying p230 (region 3 *CD2309::phoZ* reporter fusion) | This work |
| CNRS_CD386 | 630Δ*erm* carrying p197 (*CD1980.2* promoter*::phoZ* reporter fusion) | This work |
| CNRS_CD443 | CNRS_CD378 carrying p197 (*CD1980.2* promoter*::phoZ* reporter fusion) | This work |
| CNRS_CD621 | 630Δ*erm* carrying p283 (region Cdi1_3:: *phoZ* reporter fusion) | This work |
| CNRS_CD629 | CNRS_CD378 carrying p283 (region Cdi1_3:: *phoZ* reporter fusion) | This work |
| CDIP623 | CDIP601 carrying pDIA6103 (empty plasmid) | This work |
| CDIP626 | CDIP601 carrying pDIA6106 (inducible Cdi1_6-*CD1980.2* expression) | This work |
| CNRS_CD676 | 630Δ*erm* carrying pDIA6106 (inducible Cdi1_6-*CD1980.2* expression) | This work |
| CDIP1016 | CDIP601 ΔCdi1_9-*CD2309* (Δ2) | This work |
| CNRS_CD390 | CDIP1016 ΔCdi1_8-*CD1990.3* (Δ3) | This work |
| CNRS_CD523 | CNRS_CD390 ΔCdi1_10-*CD1424* ΔCdi1_11-*CD3368.2* (Δ5) | This work |
| CNRS_CD551 | CNRS_CD523 Δ*CD2386.1* Δ*CD1511.1* (Δ7) | This work |
| CNRS_CD643 | CNRS_CD551 (Δ7) carrying pDIA5987 for inducible *dccA* expression | This work |
| CNRS_CD653 | CNRS_CD551 (Δ7) carrying pDIA6103 (empty vector) | This work |
| CNRS_CD1049 | CNRS_CD551 (Δ7) carrying pDIA6106 (inducible Cdi1_6-*CD1980.2* expression) | This work |
| CNRS_CD128 | UK1 | Laboratory stock[3] |
| CNRS_CD1100 | UK1 carrying pDIA6103 (empty vector) | This work |
| CNRS_CD1101 | UK1 carrying pDIA6106 (inducible Cdi1_6-*CD1980.2* expression) | This work |
| **Plasmid** | | |

| | | |
|---|---|---|
| pMSR | Allele exchange in *C. difficile* 630 | This work |
| pDIA6103 | pRPF185 Δ*gus* vector derivative | [4] |
| pMC358 | Promoterless phoZ | [5] |
| pDIA6106 | pDIA6103 derivative for inducible Cdi1_6-*CD1980.2* expression | This work |
| pDIA6574 | pDIA6103 derivative for inducible Cdi1_6-*CD1980.2-HA* expression | This work |
| pDIA5987 | pDIA6103 derivative for inducible *dccA* expression | [6] |
| p210 | pGEMTeasy SPA-tag | Gift from E. Bentchikou |
| p197 | pMC358 derivate for fusion of promoter region of *CD1980.2,* including the Cdi1_6 riboswitch, with *phoZ* | |
| p216 | pMC358 derivate for fusion of promoter region 1 of *CD2309* with *phoZ* | This work |
| p215 | pMC358 derivate for fusion of promoter region 2 of *CD2309* with *phoZ* | This work |
| p230 | pMC358 derivate for fusion of promoter region 3 of *CD2309* with *phoZ* | This work |
| p283 | pMC358 derivate for fusion of promoter region of *flgB,* including the Cdi1_3 riboswitch, with *phoZ* | This work |

**Supplementary Table 6. Oligonucleotides used in this study.**

| Name | Sequence (5'-3')[1] | Description |
|---|---|---|
| **pDIA6103 cloning** | | |
| IMV507 | GGGATTTCTCACATAAAATAGAG | 5'pDIA6103 insert screening |
| IMV508 | TAAAATAAGCTTGATCGTAGCG | 3'pDIA6103 insert screening |
| OS499 | CCCCGAGCTCTCCTATATTATGGTAATAGTAGA | 5' Cdi1_6-*CD1980.2-SacI* |
| OS500 | CGGGATCCACTCAACTGTTTACTTTATGAA | 3' Cdi1_6-*CD1980.2-BamHI* |
| JP289 | GAACATCGTATGGGTAATTATATCTGCTTACCAATACCATC | 5' inverse PCR on pDIA6106 to add HA tag |
| JP290 | CAGATTACGCTTAAAGAAATTTTATTTTCTAGAAAACC | 3' inverse PCR on pDIA6106 to add HA tag |
| **pMSR cloning** | | |
| JP449 | CGTTTTGTAAACGAATTGC | 5'pMSR insert screening |
| JP450 | CTCACGTTAAGGGATTTTG | 3'pMSR insert screening |
| JP012 | gtttttgttaccctaagtttGGAGATAAACTATTTAATGTTGATAATTC | 5' left arm ΔCdi1_6-*CD1980.2* |
| JP013 | aatcaaattcGCTTCATTATATATTTTTCATAAAGTAAAC | 3' left arm ΔCdi1_6-*CD1980.2* |
| JP014 | taatgaagcGAATTTGATTCTTATCTAAAATGTTGG | 5' right arm ΔCdi1_6-*CD1980.2* |
| JP015 | gattatcaaaaaggagtttCCAAGTATTGCTTCTCCTATAG | 3' right arm ΔCdi1_6-*CD1980.2* |
| JP032 | GTATTATGTAAGGCTCTATACTACAATCAAG | 5' ΔCdi1_6-*CD1980.2* screening |
| JP024 | CATCAATATTCCAATTACCAATG | 3' ΔCdi1_6-*CD1980.2* screening |
| JP307 | ttttttgttaccctaagtttGTGTTTTTGTTTTTATTCATTATGG | 5' left arm ΔCdi1_9-*CD2309* |
| JP308 | ttatattaccGAACGTTAAATTATCTGAAAAAATAAAAAG | 3' left arm ΔCdi1_9-*CD2309* |

| | | |
|---|---|---|
| JP309 | tttaacgttcGGTAATATAAAATGTTTAAATTTATTTTCATATTAAG | 5’ right arm ΔCdi1_9-*CD2309* |
| JP310 | agattatcaaaaaggagtttCCATTGGTTTCTTTAAACC | 3’ right arm ΔCdi1_9-*CD2309* |
| JP314 | GCCCAAATTCATTTGTAAC | 5’ ΔCdi1_9-*CD2309* screening |
| JP312 | GTGCAGAACCTGCATAGC | 3’ ΔCdi1_9-*CD2309* screening |
| JP315 | ttttttgttaccctaagtttGCCCATATTCCATCTACAC | 5’ left arm ΔCdi1_8-*CD1990.3* |
| JP316 | aatcaatgtcCAGACTAAACACGTAGAG | 3’ left arm ΔCdi1_8-*CD1990.3* |
| JP317 | gtttagtctgGACATTGATTTCTAGAAAACC | 5’ right arm ΔCdi1_8-*CD1990.3* |
| JP318 | agattatcaaaaaggagtttGTTCCAGGAAGAAACATTAAAG | 3’ right arm ΔCdi1_8-*CD1990.3* |
| JP319 | GTAACCAACCTTTGTCATAATTACTC | 5’ ΔCdi1_8-*CD1990.3* screening |
| JP320 | GCTTCTCTGTCCATCTCTTC | 3’ ΔCdi1_8-*CD1990.3* screening |
| JP425 | ttttttgttaccctaagtttCCATCATATATTGATTCAATTTGAATATATAG | 5’ left arm ΔCdi1_10-*CD1424* |
| JP426 | gggagattagCACTAGGATAAGTTATTGTTCAAC | 3’ left arm ΔCdi1_10-*CD1424* |
| JP427 | tatcctagtgCTAATCTCCCCTATATGCAAAG | 5’ right arm ΔCdi1_10-*CD1424* |
| JP428 | agattatcaaaaaggagtttGATTGAGTTCAATTGAAAGTG | 3’ right arm ΔCdi1_10-*CD1424* |
| JP429 | CGATAATCCTCTTATAGAAACTTTAG | 5’ ΔCdi1_10-*CD1424* screening |
| JP430 | GCTGAATTTATAAGCCATAAGG | 3’ ΔCdi1_10-*CD1424* screening |
| JP431 | ttttttgttaccctaagtttGCCCACTCACCATTTTTTC | 5’ left arm ΔCdi1_11-*CD3368.2* |

| | | |
|---|---|---|
| JP432 | cttaattagcCAATAATACAATAGAATTGCTTATTATTCAATATAC | 3’ left arm ΔCdi1_11-*CD3368.2* |
| JP433 | tgtattattgGCTAATTAAGATAGATTTCTAGTATTC | 5’ right arm ΔCdi1_11-*CD3368.2* |
| JP434 | agattatcaaaaaggagtttCATATGTACCTACTGATTCC | 3’ right arm ΔCdi1_11-*CD3368.2* |
| JP435 | GTGGTATTTTGAAATTTAATGTTC | 5’ ΔCdi1_11-*CD3368.2* screening |
| JP436 | CAAATGAAGAGGTGATTTCC | 3’ ΔCdi1_11-*CD3368.2* screening |
| JP047 | cggtgtttttgttaccctaagtttCAACTAGCTGAACTTCGTTC | 5’ left arm $P_{tet}$-*CD1420* |
| JP048 | gatgcagaattcgTTTGTACTAGATTTTTGTTAAATTTTTTATTTTATTTAC | 3’ left arm $P_{tet}$-*CD1420* |
| JP049 | aatctagtacaaaCGAATTCTGCATCAAGCTAG | 5’ $P_{tet}$ module |
| JP050 | caaaataaaatacGGAGCTCAGATCTGTTAAC | 3’ $P_{tet}$ module |
| JP051 | agatctgagctccGTATTTTATTTTGGAGAAATTAATATGTTTAAAG | 5’ right arm $P_{tet}$-*CD1420* |
| JP052 | tcatgagattatcaaaaaggagtttCACAAAGGTATTTTAATACTTCATTG | 3’ right arm $P_{tet}$-*CD1420* |
| JP065 | GGAAGGTCGATTGCCC | 5’ $P_{tet}$-*CD1420* screening |
| JP064 | GAAGTGAGAAATTCCTGTTCTC | 3’ $P_{tet}$-*CD1420* screening |
| ABL22 | tttttgttaccctaagtttGGTATGTACAAGATGAAAAAG | 5’ left arm *CD2309::SPA* |
| ABL23 | atcttctcttATTATACTTACTTACCAATACCATC | 3’ left arm *CD2309::SPA* |
| ABL24 | taagtataatAAGAGAAGATGGAAAAAGAATTTC | 5’ SPA tag |
| ABL25 | aatctgtctaCTACTTGTCATCGTCATC | 3’ SPA tag |
| ABL26 | tgacaagtagTAGACAGATTTTTTTCTAGAAAAC | 5’ right arm *CD2309::SPA* |
| ABL27 | agattatcaaaaaggagtttGAAAAAATCTATTTATAATGTCATTTCC | 3’ right arm *CD2309::SPA* |
| ABL33 | CAAGTGATGGTATTGGTAAGTAAG | 5’ *CD2309::SPA* screening |
| ABL34 | GTCTAAGTTTTTGTTATGTTTTATAATG | 3’ *CD2309::SPA* screening |
| ABL78 | tttttgttaccctaagtttGGGTTTTTGTGGTTTCCTTG | 5’ left arm *CD2309-spoZ* |
| ABL79 | tacattgacgCTAAAAAACTTTCATGAAAAACAAAAAG | 3’ left arm *CD2309-spoZ* |

| | | |
|---|---|---|
| ABL80 | agtttttagcGTCAATGTATGGGTAGATATG | 5’ RBS-*spoZ* |
| ABL81 | aaaaaatctgGTTATTTGCCAATACCTTTATC | 3’ RBS-*spoZ* |
| ABL82 | ggcaaataacCAGATTTTTTTCTAGAAAACCC | 5’ right arm *CD2309-spoZ* |
| ABL83 | agattatcaaaaaggagtttCAAATTAACTTATTGTAGATTTTTTATTTTTTAAC | 3’ right arm *CD2309-spoZ* |
| ABL84 | CTAAGGAGGATAAAAAATGATTCAG | 5’ *CD2309-spoZ* screening |
| ABL85 | CTAAAAAGGGTTTTCTAGAAAAAAATC | 3’ *CD2309-spoZ* screening |
| **pMC358 cloning** | | |
| JP725 | CGTCAATGTATGGGTAGATATG | 5’ pMC358 linearization for construction of transcriptional fusions |
| JP726 | CAACGTCGTGACTGGG | 3’ pMC358 linearization for construction of transcriptional and traanslational fusions |
| ABL01 | AAGAAAAGAGCTTTGCTAGG | 5’ pMC358 linearization for construction of translational fusions (removal of *phoZ* RBS) |
| ABL02 | ttttcccagtcacgacgttgGGATGTATATCAATTCCATACAAATATTG | 5’ Cdi1_6 region |
| ABL03 | tatctacccatacattgacgCTAAAAAACATTCATAAAAACAAAAAG | 3’ Cdi1_6 region |
| ABL07 | ttttcccagtcacgacgttggCTATAAAACTTTATATTTTAGAGGAAAAATAATATG | 5’ Cdi1_9 region |
| ABL08 | tatctacccatacattgacgCTAAAAAACTTTCATGAAAAACAAAAAG | 3’ Cdi1_9 region 2 |
| ABL09 | tatctacccatacattgacgcGATTTAGTTTTATTCTACAGATTAATAATAC | 3’ Cdi1_9 region 1 |
| ABL010 | cctagcaaagctcttttcttCATAATGTATTTATCCCCCTAAAAAAC | 3’ Cdi1_9 region 3 |
| ABL035 | ttttcccagtcacgacgttgGTTACTCTATACAACATTTTAGAC | 5’ Cdi1_3 region |
| ABL036 | tatctacccatacattgacgGTAAAATAGTAACTTTTTGAGATAATTTTTC | 3’ Cdi1_3 region |
| JP733 | CTATTACGCCAGCTGGC | 5’pMC358-screening |
| JP734 | GGTAACCCCTAGCAAAGC | 3’pMC358-screening |
| **qRT-PCR** | | |

| | | |
|---|---|---|
| polIIIF | TCCATCTATTGCAGGGTGGT | 5’ *DNA pol III* |
| polIIIR | CCCAACTCTTCGCTAAGCAC | 3’ *DNA pol III* |
| OS576 | TGGATAAATGTACACATATGTTAACTGC | 5’ *CD2309* |
| OS577 | TGCAAGCAAGATTCCACAAC | 3’ *CD2309* |
| JP335 | GTTCACGCGATTACTGC | 5’ *CD1980.2* |
| JP336 | TCTGCTTACCAATACCATCA | 3’ *CD1980.2* |
| OS587 | GCAGTTCGTATGATTATTGTAATTTTT | 5’ *CD1990.3* |
| OS588 | CCATTACTTGGTGTAAACAAGACTC | 3’ *CD1990.3* |
| JP343 | GATGTACACATATGTTGACTGC | 5’ *CD3368.2* |
| JP344 | AAACAAGATTCTACCACATTTTC | 3’ *CD3368.2* |
| JP337 | CTATTCATTCATAGAAACTCAAATTG | 5’ *CD1511.1* |
| JP338 | CAAACCACATTCGCATAGT | 3’ *CD1511.1* |
| JP339 | GTGGATATTATTCATTTATTGACAC | 5’ *CD2386.1* |
| JP340 | CATTTACAAAGCAACATCATTATAC | 3’ *CD2386.1* |
| JP947 | GGCCAAAGTGTAATGCAAGG | 5’ *CD2831* |
| JP948 | GCATCTGGAACATCCGTTTT | 3’ *CD2831* |
| OS580 | CAGTAGTGGCAGTTCCAGCTT | 5’ *pilA1* |
| OS581 | CCAGTTTGACCATCTGGTGT | 3’ *pilA1* |
| OS574 | TTCCTAAGGGATGGGAAGGT | 5’ *ppep-1* |
| OS575 | ATTGCATCATGCCCTTTACC | 3’ *ppep-1* |

[1]Underlined bases indicate engineered restriction sites; lowercase bases indicate overlapping sequences
